# Decoding microbial diversity in roots of rice plants under flooded conditions: influence of the host genotype, root compartment and mycorrhizal association

**DOI:** 10.64898/2026.01.22.701102

**Authors:** Iratxe Busturia, Héctor Martín-Cardoso, Concha Domingo, Antoni Garcia-Molina, Blanca San Segundo

## Abstract

**Background:** The root microbiome plays a critical role in nutrient acquisition, stress tolerance and overall plant health. Rice, a staple food for more than half of the world’s population, is commonly cultivated under flooded conditions. Despite its agronomical importance, our current understanding of root-associated microbiomes in rice grown under flooded conditions is limited. On the other hand, nitrogen (N) and phosphorus (P) fertilizers are routinely applied to maximize rice yield. It is also well known that root colonization by arbuscular mycorrhizal (AM) fungi enhances mineral nutrition in plants, but whether mycorrhizal associations influence the composition of the rice root microbiome remains poorly understood. In this study, shotgun metagenomic sequencing was used to characterize the root endosphere and rhizosphere microbiomes in two temperate *japonica* rice varieties (cv. Bomba and JSendra) grown under flooded conditions. The impact of colonization by the AM fungus *Rhizophagus irregularis* on the root microbiome was investigated.

**Results:** Root-associated compartments harbour distinct microbial communities in rice with bacterial taxa comprising approximately 95% of the total microbia in rice roots. At the Phylum level, the root bacteriome was primarily composed of Pseudomonadota (Alphaproteobacteria, Betaproteobacteria and Gammaproteobacteria) followed by Actinomycetota. The fungal microbiome was dominated by Ascomycota (Sordariomycetes, Eurotiomycetes and Dothideomycetes) and Basidiomycota. Not only the root compartment, but also the host genotype can shape the root microbiome. Recruitment of specific microorganism mainly occurs at the species level. Genotype-specific and compartment-specific associations of microbial species in mycorrhizal rice roots were also observed supporting that root colonization by an AM fungus contributes to variations in the root microbiome. Further, key microbial species primarily associated to methane production and nutrient cycling (e.g. Phosphate Solubilizing Bacteria and Nitrogen cycling bacteria) colonizing root compartments in each rice genotype and mycorrhizal condition are described.

**Conclusions:** The rice genotype, root compartment and mycorrhizal condition markedly influence the microbiome in roots of rice plants growing in flooded rice fields. These findings illustrate the potential of the plant to shape its associated root microbiome, thus, offering valuable insights for the development of microbiome-based strategies to improve growth and performance in rice plants under flooded conditions.

## Introduction

In natural and agricultural environments, plants and soil microbes engage in complex dynamic interactions that might have different effects on the host plant. These associations can be beneficial (e.g., by facilitating nutrient uptake or promoting resistance to environmental stresses) or might have adverse effects (e.g., by promoting disease or outcompeting the plant for resources) on the host plant [1]. In this way, plants cannot longer be viewed as autonomous entities but rather as a more complex entities composed of the host plus the associated microbes.

Rice serves as a crucial staple food for a large part of the world’s population. To meet the requirements of the increasing global population, there is a growing need to increase rice production [2]. Over the past decades, there has been an enormous increase in the use of fertilizers and pesticides to optimize yield, which is also the principal cause of pollution in major rice-producing regions. The overuse of fertilizers also contributes to soil degradation, eutrophication in aquatic ecosystems, loss of biodiversity, and imbalance in microbial community structures. Harnessing the plant microbiome offers new opportunities to improve plant nutrition and health for the development of sustainable rice cultivation systems [3–5].

Rice, unlike other cereals, is usually grown under flooded conditions (or paddy fields), As the rice plant must select microorganisms inhabiting the flooded soil to assemble its own root microbiome, the composition and functional features of the rice root microbiome is greatly influenced by flooding [6]. Indeed, rice paddies creates a unique anaerobic environment ideal for specific microorganisms, such as methanogenic archaea. On the other hand, cycling of nutrients, e.g., carbon (C), nitrogen (N) and phosphorus (P), represents an important part of the ecosystem function in agricultural soils. Hence, microbial communities involved either in nutrient acquisition by plant roots or nutrient cycling are particularly important for healthy plant growth in paddy fields [7].

Arbuscular mycorrhizal (AM) fungi enhance plant uptake of nutrients (mainly phosphorus and nitrogen) from the soil and the transferring to the plant. The improved nutrient acquisition due to AM symbiosis might also have a great impact on plant growth and productivity, an aspect that remains largely unexplored in flooded rice [5]. Evidence also support that Phosphate-Solubilizing Bacteria (PSB) improve plant nutrition by converting insoluble P in soils into usable P that can be directly taken by plants. Ultimately, interactions between rice roots and soil microbes, and microbe-microbe interactions would determine plant performance and microbial ecological functions in paddy field ecosystems.

Efforts carried out so far to decipher microbiome assemblages in different rice tissues (e.g., leaves, roots, seeds) demonstrated a remarkable diversity and dynamics in rice microbiomes [5, 6, 8–14]. Microbial communities in root-associated compartments, namely the root endosphere and rhizosphere compartments, might also vary depending on the geographical location, soil type, management practices, environmental factors, and/or host genotypes [3, 15]. Remarkably, most studies so far carried out to investigate root-related microbiomes in rice have been conducted on plants grown under aerobic conditions, while few studies have been conducted in rice plants growing in paddy fields. Understanding microbial diversity and functions of microbial communities in flooded rice ecosystems is a requisite for the development of effective sustainable agricultural practices in rice cultivation.

In recent years, the availability of hight-throughput sequencing and metagenomic techniques has accelerated studies on plant microbiomes. Most of these studies on the characterization of rice microbiomes used metabarcoding amplicon sequencing (16S rRNA gene) [9–12, 16, 17]. Unlike amplicon sequencing, shotgun metagenomics captures total genomic DNA without target-specific PCR amplification, thereby providing a more comprehensive information on the entire microbial community’s taxonomic composition.

Although numerous microbes inhabit the soil in paddy fields, only a part of them will inhabit the root rhizosphere, and only some of these microbes would enter the root endosphere. Studies on the root-associated microbiomes of rice plants grown under aerial seedling cultivation systems revealed a greater proportion of *Proteobacteria* and *Spirochaetes* in the endosphere than the rhizosphere [16]. Differences in the abundance of Beta- and Gamma-proteobacteria (phylum *Pseudomonadota*) between the rhizosphere and root endosphere were observed in rice varieties grown in different geographical locations in China, with Gamma-proteobacteria being the most dominant taxa in the root endosphere of those varieties [11]. Presumably, root-associated microbial communities might have been selected by flooded rice cultivation. Deciphering differences in microbial communities in the root endosphere-rhizosphere microbiomes in rice plants growing under flooded conditions will help in developing strategies to enhance health and productivity in rice cultivation. At present, however, we are only at the beginning of understanding the structure of root-associated microbiomes in paddy fields.

To address this gap of knowledge in rice, in this study we addressed the root endosphere and rhizosphere microbiomes in two temperate *japonica* rice varieties of agronomical interest, Bomba and JSendra, grown in paddy fields. Recently, we described that the two rice varieties here investigated can be efficiently colonized by *Rhizophagus irregularis* [18, 19]. We therefore studied the effect of inoculation with the AM fungus *R. irregularis* on microbial communities colonizing the root endosphere and rhizosphere in Bomba and JSendra plants in paddy fields. Analysis of metagenomic datasets allowed the identification of microbial taxa involved in P solubilization and N cycling in each compartment and condition in Bomba and JSendra plants. Results obtained in this work demonstrate that the root compartment (endosphere and rhizosphere), genotype and mycorrhizal status of the rice plant are the key factors shaping the rice root-related microbiomes in flooded rice systems.

## Materials and methods

### Rice genotypes, experimental design, and sample collection

Two commercial rice cultivars (*O. sativa* cv *japonica*) Bomba and JSendra were used in this study. Seedlings were grown on seedling beds in a nursery under aerobic conditions at 28°C, 14h/10h light/dark cycle and high humidity as previously described [19]. Half of the plants were inoculated with the AM fungus *Rhizophagus irregularis* (MycAgro, Bretenière, France, http://www.mycagrolab.com) while the other half was mock-inoculated (no AM fungi added to the soil) [19]. Five weeks after AM fungus inoculation, the rice seedlings were transplanted into flooded rice fields. As previously reported, 5 weeks of aerobic growth after inoculation with the AM fungus was enough time to ensure penetration of the fungus into the rice roots and subsequent establishment of the AM symbiosis in rice plants growing into flooded rice fields [19]. The two rice cultivars were grown in fields located in a traditional rice producing region of eastern Spain (Valencia, 39°15^..^46.9”N 0°20^..^41.2”W). Mock-inoculated and *R. irregularis*-inoculated plants were grown until maturity (May-September, 2024). Three plots (each plot containing 30 plants) were established in the same field for each variety (Bomba, JSendra) and condition (*R. irregularis* and mock-inoculated), maintaining a distance of 4 m between non-mycorrhizal and mycorrhizal plots (**Additional file 1: Fig. S1**). The experimental field was exclusively used for rice cultivation. Neither fertilizers nor pesticides were applied to the rice plants in the field.

Samples were collected at the harvest time from two niches: root endosphere and rhizosphere. For metagenomic analysis of the root endosphere, three samples from each experimental plot were examined (each sample was derived from a pool of roots from two rice plants). The corresponding rhizosphere samples were pooled and examined.

For metagenomic analysis, a total of 36 endosphere samples were examined (2 genotypes x 2 conditions x 3 biological replicates x 3 plots). The corresponding 36 rhizosphere samples were also used for metagenomic analysis. Rhizosphere samples were collected from the most adjacent soil surrounding the roots. For endosphere samples, five washings steps were carried out to discard soil sediments in rice roots. Rhizosphere and root endosphere samples were pulverized before DNA extraction.

### DNA extraction

DNA from rice roots was extracted using MATAB as the extraction buffer (0.1 M of Tris–HCl pH 8.0, 1.4 M NaCl, 20 mM EDTA, 2% MATAB, containing 1% PEG 6000 and 0.5% sodium sulfite) according to [20]. Extraction of DNA from rhizosphere samples was carried out using the NucleoSpin Soil kit (Macherey-Nagel, Düren, Germany).

### Metagenome Sequencing

Shotgun sequencing was used to examine the root-associated microbiomes; endosphere and rhizosphere. For library preparation, the genomic DNA was randomly fragmented by sonication, and the obtained fragments were then end-repaired, A-tailed, and ligated with Illumina adapters, followed by PCR amplification using P5 and P7 primers (P5, 5’-AGATCGGAAGAGCGTCGTGTAGGGAAAGAGTGTAGATCTCGGTGGTCGCCGTATCATT-3’; P7, 5’-GATCGGAAGAGCACACGTCTGAACTCCAGTCACGGATGACTATCTCGTATGCCGTCTTCTGCTTG-3’). The resulting PCR products were size selected, PCR amplified and purified with an AMPure XP system. The libraries were examined for size distribution and quantified through Qubit and real-time PCR. Quantified libraries were sequenced (2 x 150 bp paired-end sequencing) on the Illumina platform at Novogene (Cambridge, UK) (depth of 6 Gb/library).

### Bioinformatic and Statistical analysis

Quality check of raw sequencing data was performed using FastQC *v.0.12.1* [21], and adaptor sequences were trimmed with fastp *v.0.24.0* (Q = 20, minimum read length = 30) [22]. Trimmed and filtered reads were mapped to the *Oryza sativa* var. Nipponbare reference genome IRGSP-1.0 (https://plants.ensembl.org/Oryza_sativa/Info/Index) using Bowtie 2 (*v.2.5.4*) [23]. Positive hits with the *O. sativa* genome were removed to produce a final set of clean sequencing data. Metagenomes were assembled independently for each sample. The *R. irregularis* genome assembly ASM43914v3 (https://fungi.ensembl.org/Rhizophagus_irregularis_daom_181602_gca_000439145/Info/Index) was used to identify reads from the AM fungus in the various metagenomic datasets.

Taxonomy assignment of metagenomic sequencing reads was done with Kraken2 *v2.1.5* software and the Kraken core nt database [24]. Bracken *v.3.1* (Bayesian Reestimation of Abundance with KrakEN) was used to estimate the relative abundances of microbial taxa at the species level [25]. Reads classified as bacterial and fungal reads were analyzed in R *v.4.3.3* with functions of R packages (phyloseq *v.1.52.0*, ggplot2 *v.4.0.0*, tidyverse *v.2.0.0,* vegan *v.2.7-1,* microbiome *v.1.31.3* and pairwise Adonis *v.0.4.1*) [26–30].

Principal Coordinates Analysis (PCoA) was performed using the functions from the phyloseq R and microbiome packages using Bray-Curtis distances [27]. Homogeneity of dispersion between groups of samples was tested with *betadisper* and a non-parametric test for significance with *permutest* functions. A Permutational Multivariate Analysis of Variance (PERMANOVA) based on Bray-Curtis distances using *adonis2* function was performed to evaluate statistically differences between groups.

Shannon measurements were used to evaluate α-diversity indices in the bacterial and fungal communities across samples. Shannon measurements not only reflect the number of species in the sample (richness) but also reflect the abundance distribution of each species in the sample (evenness) [31, 32]. Differences between samples were tested with a Kruskal Test followed by a *post-hoc* analysis with Dunn Test. Hierarchical clustering heatmaps were used to illustrate differences in abundance of species across the different samples. Here, a log(x+1) transformation was done, followed by the Min–Max Normalization (MMN) method [33]. Sankey diagrams were produced using the Pavian R package *v1.2.1* [34].

Differential abundance analysis of microbiome data was carried out at species level by ANCOM-BC2 *v.2.10.0* (Analysis of Compositions of Microbiomes with Bias Correction 2) [35]. Changes in relative abundance between two groups of samples with biological consistency were determined using adjusted p-values with multiple testing correction according to Holm-Bonferroni method (adjusted *P* D 0.05).

## Results

### Overall microbial community composition in root-associated compartments of rice plants

The structure and diversity of microbial communities in the root endosphere and rhizosphere of the rice varieties Bomba and JSendra (*japonica* rice cultivars) were investigated in plants cultivated under flooded conditions. To minimize the risk of variations in environmental or geographical factors potentially affecting the root microbiome (e.g., soil type, management practices), the plants were grown in the same paddy field, located in a traditional rice-growing area in Valencia (eastern Spain). Additionally, knowing that the AM symbiosis increases yield and disease resistance in both Bomba and JSendra plants [19], the impact of inoculation with the AM fungus *R. irregularis* in the root microbiome of both rice varieties plants was also addressed. Methodologically, shotgun metagenomics was employed to provide broader coverage and higher resolution in taxonomic profiling of metagenomic data compared to amplicon sequencing.

A total dataset of 272.84 Gb was generated from endosphere samples (3 experimental plots x 3 replicates each plot x 2 genotypes x 2 conditions; 36 samples), plus rhizosphere samples (3 plots x 3 replicates each plot x 2 genotypes x 2 conditions; 36 samples). After trimming the reads, a total of 197 Gb (1,313.29 million, 150 bp-end reads) passed the quality control parameters (**Additional file 2: Table S1**). Removal of host reads yielded a final set of clean reads from the different samples which were assembled independently (1,313,291,149 clean reads in total). To confirm AM colonization, metagenomic reads were mapped to the reference genome of *R. irregularis* ASM43914v3, revealing an average of 13,362.5 and 12,631.39 reads - the 0.07% and 0.05% of the total reads in the endosphere – derived from this fungus in AM-inoculated Bomba and JSendra plants, respectively (**Additional file 2: Table S1**). In the non-inoculated plants, *R. irregularis* reads lowered to 0.043% and 0.046 % of the total number of reads (in endosphere) in Bomba and JSendra, respectively (7,436.44 and 10,532.94 reads) (**Additional file 2: Table S1**).

Taxonomic classification of metagenomic datasets was carried out using Kraken2 (v2.1.5). Bacterial taxa greatly exceeded fungal taxa across samples (**Additional file 1: Fig. S2**). On average, 94-96% of the total microbia corresponded to Bacteria, while fungal taxa represented 2%-3% of the microbial community. The complete list of species identified in each metagenomic dataset is presented in **Additional file 2: Table S2**.

Principal coordinate analysis (PCoA) with Bray-Curtis distances was used for unsupervised examination of microbial beta diversity to verify similarity and dissimilarity in microbial community structure between different samples. The principal coordinate 1 (PCo1, explaining 82.7-80.9% of total variance in bacterial and fungal communities, respectively) mainly separated root endosphere from rhizosphere communities, regardless of the mycorrhizal status (**Fig. 1A**). Indeed, permutation multivariate analysis of variance (PERMANOVA) test corroborated significant differences between the endosphere and the rhizosphere compartments (*P* = 0.001; F = 51.377 for bacteria; F = 21.241 for fungi). (**Fig. 1A**). In a second term, the composition of bacterial communities could be further discriminated by the rice genotype at the level of root endosphere (PCo2 7.8% total variance), but not in the rhizosphere microbiome (**Fig. 1A**). Notably, the bacterial communities within the endosphere of AM-inoculated plants were slightly discriminated from those not colonised (**Fig. 1A**). Thus, our primary analysis points to important different plant microbiome composition depending on the compartment (rhizosphere vs endosphere) and, within endospheres, due to the rice variety and, to certain extend, to AM-colonisation. Consequently, subsequent analyses in AM and non-AM-inoculated plants were conducted separately for each genotype and compartment.

**Fig. 1.**
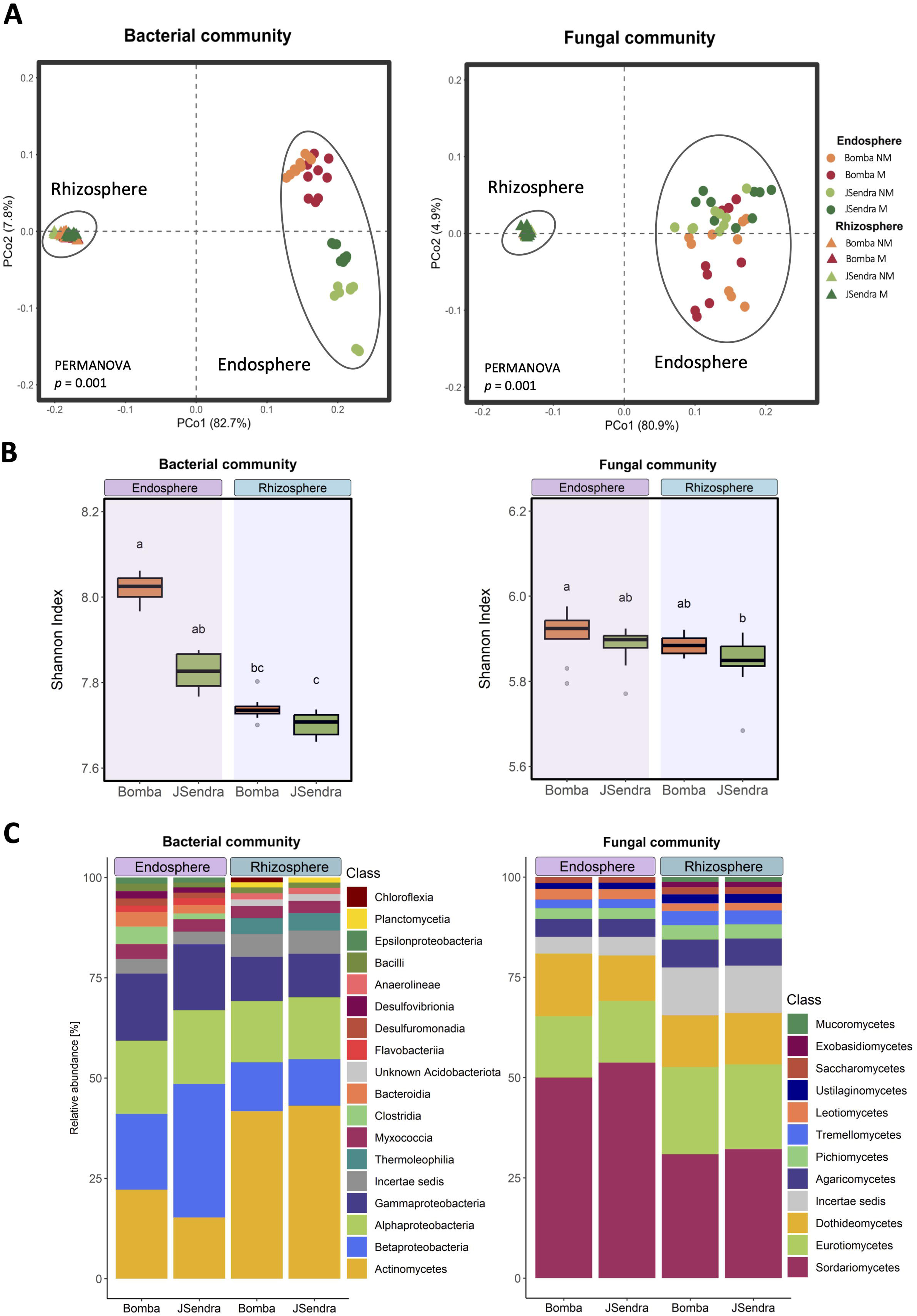
Influence of the root compartment and rice genotype on bacterial and fungal communities in rice. Plants (cv. Bomba and JSendra) were grown in rice fields under flooded conditions as previously described [19]. For each variety, two sets of plants were grown in parallel. One set of plants was pre-inoculated with the AM fungus *R. irregularis* before transplanting to the rice field, whereas the other set of plants was mock-inoculated [19]. Roots were collected at the harvest time. For each compartment, the bacterial and fungal microbiomes, were examined separately. **(A)** Principal coordinates analysis (PCoA) of beta-diversity, measured by Bray-Curtis distances of bacterial (left panel) and fungal (right panel) communities in the endosphere (circles) and rhizosphere (triangles) of Bomba and JSendra plants. Each PCoA plot indicates the percentage of variability explained by each factor based on PERMANOVA. The light and dark colour represents the mycorrhizal condition: NM, non-mycorrhizal; M, mycorrhizal condition. Ellipses added manually for visual emphasis. (**B**) Alpha Shannon diversity of bacterial and fungal communities in the root endosphere and rhizosphere of rice (cv. Bomba and JSendra) plants, in non-mycorrhizal plants. Different letters indicate statistically significant differences among samples (Dunn’s Test, P D 0.05). (**C)** Relative abundance of microbial taxa at the Class level in the root endosphere and soil rhizosphere of Bomba and JSendra plants (non-mycorrhizal condition). Taxa that exhibit a relative abundance > 1% are presented.

### Comparative study of the root endosphere and rhizosphere microbiomes in rice varieties

To gain more insight into the microbiome of compartments and genotypes, the Shannon index was used to estimate the differential composition in bacterial and fungal communities by considering both richness (number of species) and evenness (variability in species abundances) within samples. The bacterial community in the endosphere of non-AM Bomba and JSendra displayed significantly higher values of Shannon indexes compared to the rhizosphere, indicating higher species richness and evenness (**Fig. 1B**, left panel). Within the endosphere, the bacterial community in Bomba was more diverse than in JSendra, supporting distinct capacity of rice genotypes to shape the bacterial community at this root level (**Fig. 1B**, left panel). Shannon diversity indexes indicated minimal differences between genotypes in the rhizosphere, revealing a high similarity in the bacterial community colonizing this compartment (**Fig. 1B**, left panel). Conversely, neither the compartment, nor the rice genotype could influence the fungal community residing in the root endosphere – Shannon indexes barely altered (**Fig. 1B**, right panel).

Next, the distribution of taxa in our metagenomic datasets was assessed per compartment and genotype. Among the most abundant taxa, the most prevalent Classes in the bacterial community across the two compartments were Actinomycetes, Betaproteobacteria, Alphaproteobacteria and Gammaproteobacteria (**Fig. 1C**, left panel). However, the proportions varied among genotypes and compartments. For instance, the proportion of Actinomycetes at the endosphere was the half of the ones at the rhizosphere in both genotypes, while Betaproteobacteria were 1.5- to 2.5-fold more abundant in the endosphere compartment (**Fig. 1C**, left panel). In the fungal community, the dominant taxa at the Class level in all the samples were Sordariomycetes followed by Eurotiomycetes and Dothideomycetes, their relative abundance also varying among compartments (**Fig. 1C**, right panel). Of them, the representation of Sordariomycetes was the half in the rhizosphere than in the root endosphere in both rice varieties, while Eurotiomycetes were more abundant in the rhizosphere compartment (**Fig. 1C**, right panel). Between species, differences in abundances of bacterial and fungal species in the endosphere and rhizosphere of rice varieties were observed (**Additional file 1: Fig. S3**). For both bacteria and fungi, a cluster specific for rhizosphere can be observed. On the other hand, the endosphere of both communities presents not only a cluster with common species between varieties, but also a cluster with variety-specific species. Then, our analysis supported that, although the compartment type is major factor shaping the composition of microbial species in rice roots, there are also genotype-specific associations of microbial species.

For a better understanding of microbial diversity in root-associated compartments of Bomba and JSendra plants, subsequent analyses were carried out at the species level. Sankey were elaborated with the top 20 most abundant species to visualize differences among taxons, bacterial and fungal taxons, colonizing each root-compartment of Bomba and JSendra roots. Considering the bacterial microbiome, Pseudomonadota and Actinomycetota were the most abundant phyla in the endosphere of the two rice genotypes, followed by Bacillota and Bacteroidota (**Fig. 2**, **Additional file 2: Table S3**). Pseudomonadota accounted for approximately 46.59% and 60.83% of all bacteria in the endosphere of Bomba and JSendra, respectively. In the endosphere of JSendra, Betaproteobacteria, Alphaproteobacteria and Gammaproteobacteria accounted for 29.58%, 16.35% and 14.66 % of all microbia, while these Classes showed a similar abundance in the endosphere of Bomba plants (16.25 %, 15.7% and 14.42%, respectively) (**Additional file 2: Table S3**). In particular, we noticed that *Hydrogenophaga* species (Class Betaproteobacteria) were among the most abundant species in the endosphere of JSendra, but not in Bomba (**Fig. 2**). *Hydrogenophaga* consist of bacteria capable of using hydrogen as an energy source and are related to hydrogen autotrophic denitrification [36]. On the other hand, *Sulfurospirillum* sp. (Class Epsilonproteobacteria) was identified among the most abundant species in the endosphere of Bomba, but not in JSendra (**Fig. 2**). Then, it will be of interest to investigate whether root colonization by *Sulfurospirillum* provides an advantage to Bomba plants in growing in contaminated fields. Bacterial species in the Bacillota and Bacteroidota phyla were less represented in the endosphere of Bomba roots than in JSendra roots (**Fig. 2**, **Additional file 2: Table S2**).

**Fig. 2.**
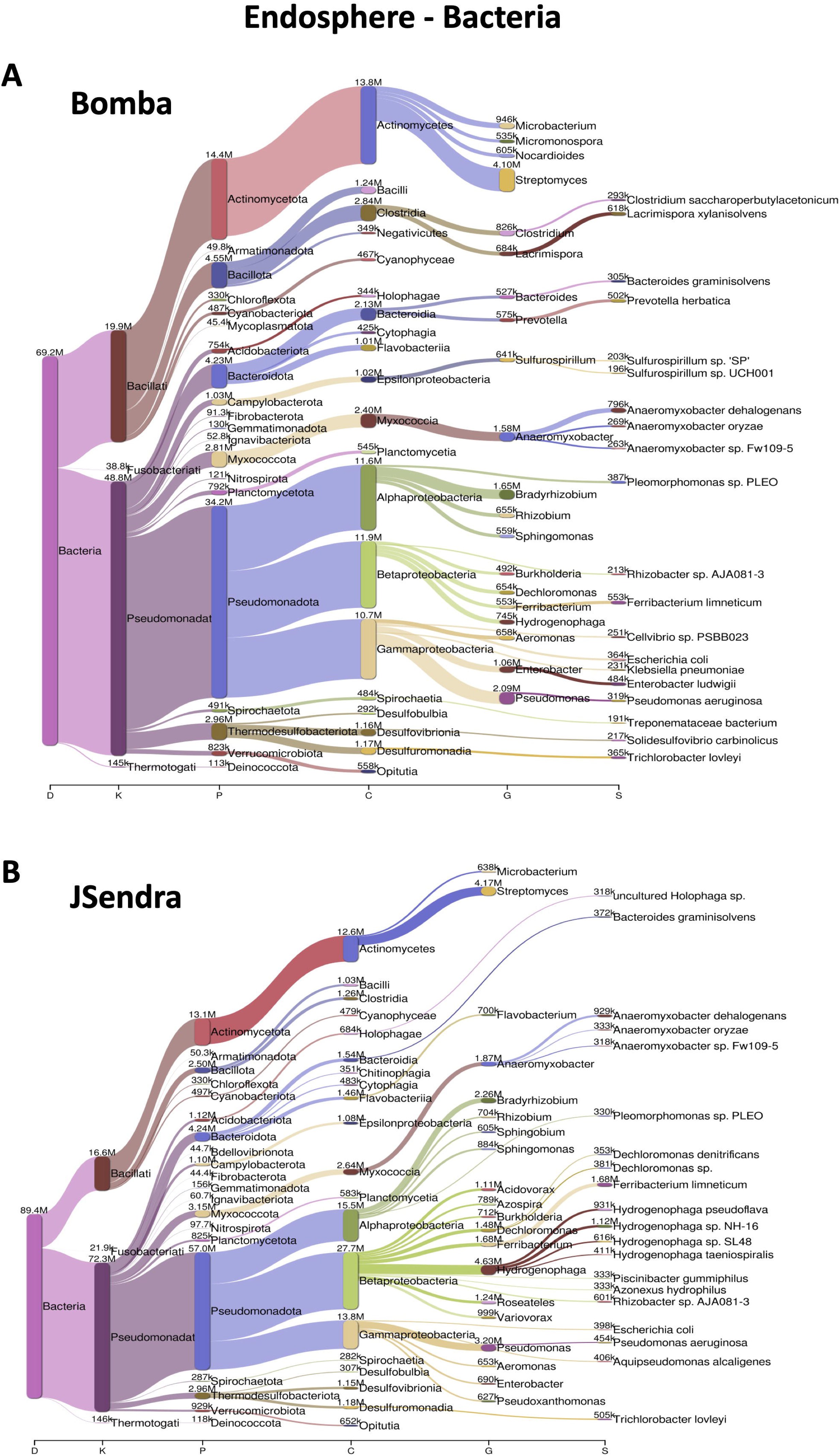
Most abundant bacterial taxa identified in the root endosphere of rice plants. Sankey diagrams for Bomba (**A**) and JSendra (**B**) plants are presented. The relative abundance of the top 20 taxa in each variety is presented, from the domain (on the left) to species (on the right). The width of the bar is proportional to the abundance, whereas nodes depict the hierarchy levels of microbial taxa.

Regarding the fungal microbiome colonizing the endosphere of Bomba and JSendra roots, the most predominant fungal phyla in the endosphere were Ascomycota (80.5% and 80.15% of all fungi for both Bomba and JSendra, respectively), followed by Basidiomycota (9.31% and 9.57% of all fungi for both Bomba and JSendra, respectively) (**Fig. 3**, **Additional file 2: Table S3**). Fungal species identified in the endosphere of rice varieties included potential pathogenic fungi for rice. For instance, *Alternaria alternata* was among the most abundant fungal taxa in the endosphere of Bomba roots, but not in the endosphere of JSendra roots (**Fig. 3**). *Fusarium oxysporum* and *Nigrospora oryzae* were identified in the endosphere of both varieties (**Fig. 3**). *Thermothelomyces* and *Thermothielavioides* species were also among the most abundant fungal species in the endosphere in both rice varieties. Compared with the endospheric bacterial microbiome, the host genotype appears to have a weaker effect on the fungal community inhabiting the root endosphere in rice varieties investigated in this study.

**Fig. 3.**
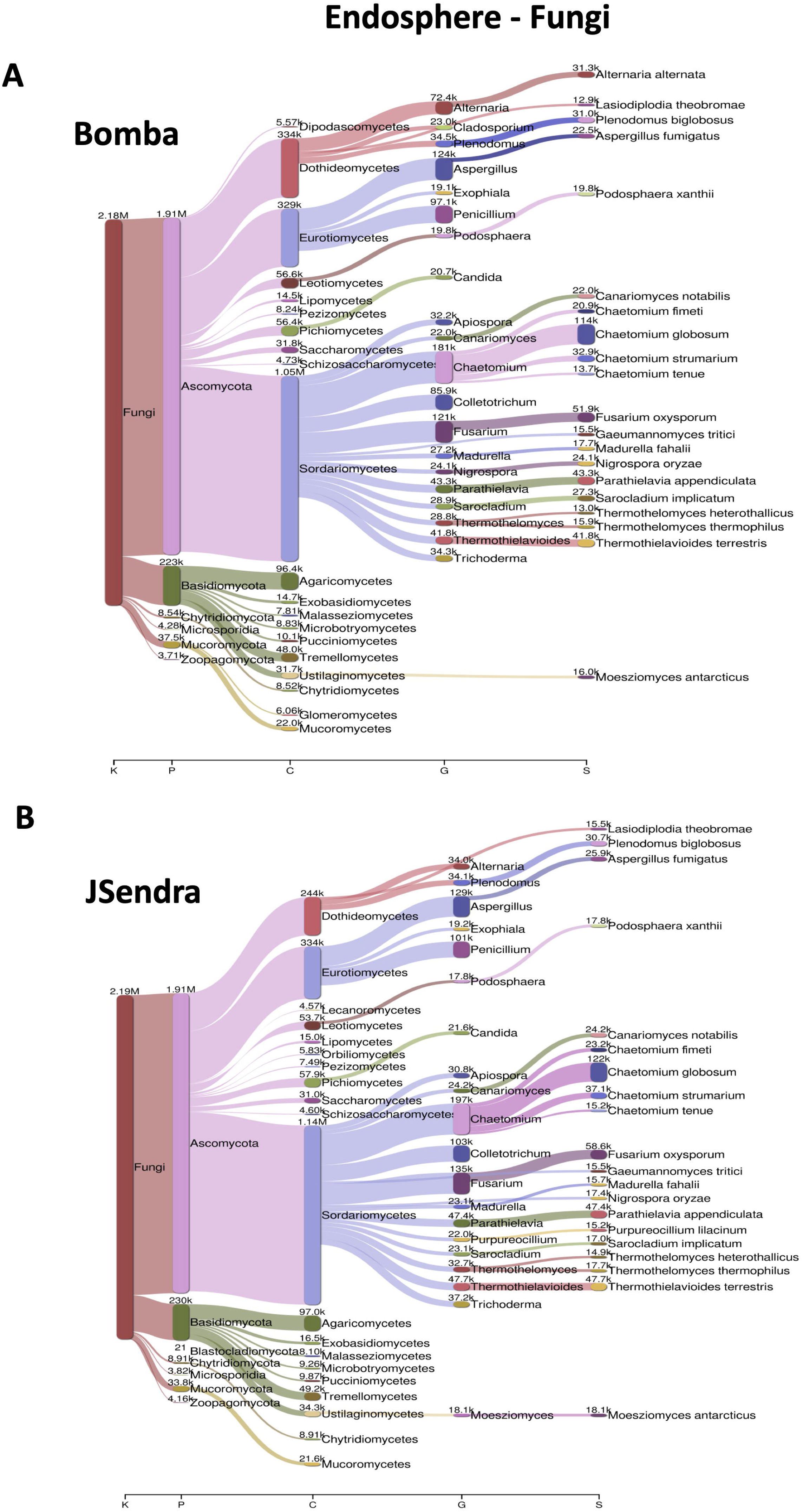
Most abundant fungal taxa identified in the root endosphere of rice plants. Sankey diagrams for Bomba (**A**) and JSendra (**B**) plants are presented. The relative abundance of the top 20 in each variety is presented, from the kingdom (on the left) to species (on the right). The width of the bar is proportional to the abundance, whereas nodes depict the hierarchy levels of microbial taxa.

Finally, we examined the bacterial and fungal communities in the rhizosphere of rice varieties. The bacterial community was dominated by taxa in the Phyla of Pseudomonadota (accounting for 33.57% and 32.92% of bacteria in Bomba and JSendra plants, respectively), and Actinomycetota (41.51% and 42.75% of bacteria in Bomba and JSendra, respectively) (**Fig. 4**, **Additional file 2: Table S3**). At the species level, distinct taxa were specifically enriched in the rhizosphere of one or another rice variety. For instance, the plant growth-promoting bacterium *Delftia acidovorans* (Phylum Pseudomonadota) was identified among the most abundant species in the rhizosphere of Bomba, but not in the rhizosphere of JSendra. *D. acidovorans* has been reported to have the potential of functioning in bioremediation in rice paddies contaminated with heavy metals like cadmium [37]. The rhizosphere fungal community was dominated by Ascomycota in both rice varieties, followed by Basidiomycota (**Additional file 1: Fig. S4**). However, no major differences were detected in the taxonomic profile of the most abundant fungal species in the rhizosphere of rice varieties (**Additional file 1: Fig. S4**).

**Fig. 4.**
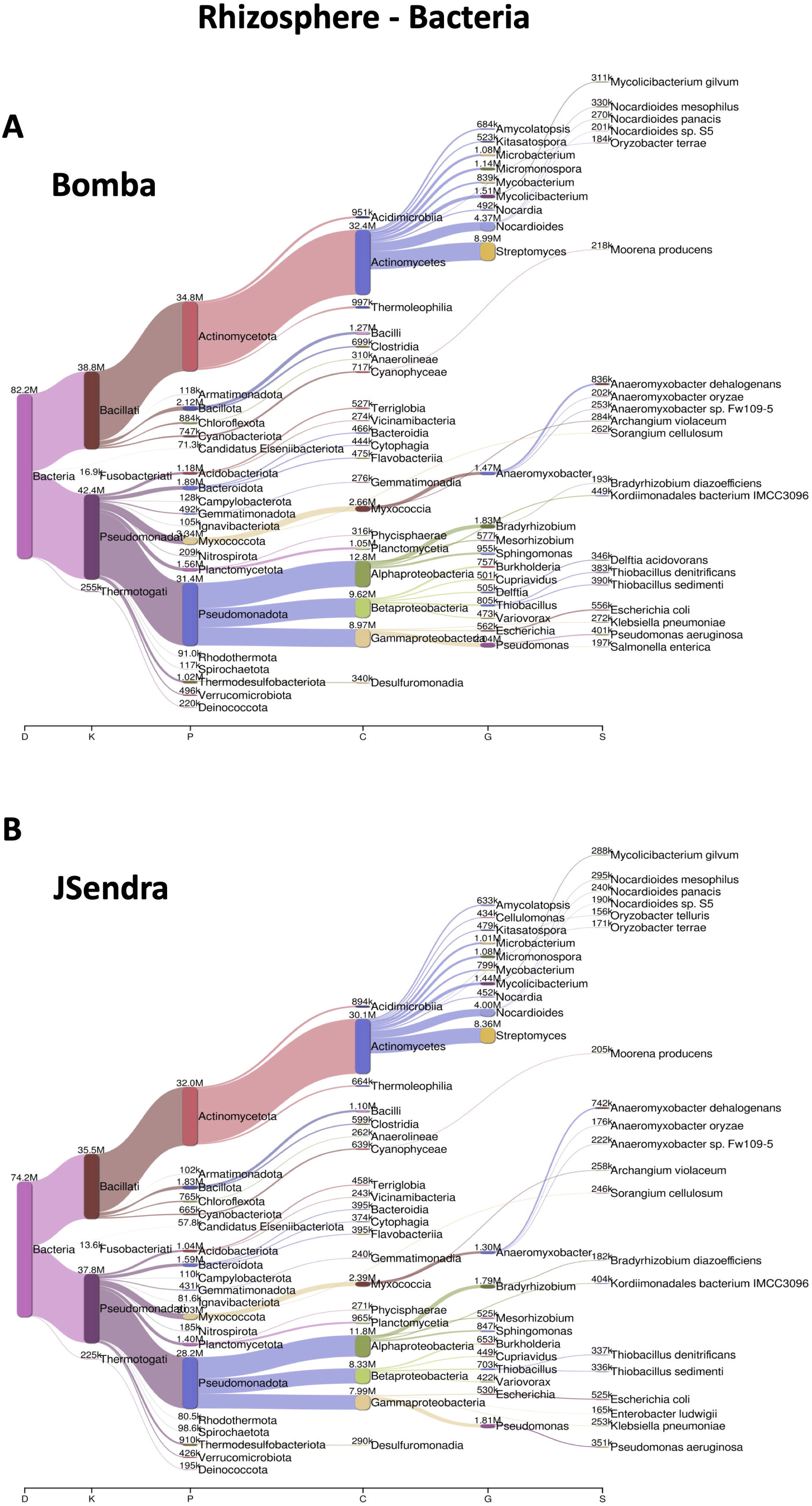
Most abundant bacterial taxa identified in the rhizosphere of rice plants. Sankey diagrams for Bomba (**A**) and JSendra (**B**) plants are shown. The relative abundance of the top 20 taxa in each variety is presented, from the kingdom (on the left) to species (on the right). The width of the bar is proportional to the abundance, whereas nodes depict the hierarchy levels of microbial taxa.

Overall, these analyses revealed species-level variations in microbial assemblages, mainly in bacterial community structures, between root-associated compartments of rice, which were also dependent on the rice genotype. Differences observed in abundance and composition of microbial communities between the endosphere and rhizosphere compartments support that not all microbes inhabiting the rhizosphere are capable of colonizing the rice root endosphere, while pointing to a preferential recruitment of microbes from the rhizosphere in rice roots.

### The arbuscular mycorrhizal symbiosis shapes the bacterial and fungal community in rice

Previous studies revealed that inoculation of rice plants, especially Bomba and JSendra, with the AM fungus *R. irregularis* increased productivity and enhanced disease resistance [18, 19]. However, it is unclear whether root colonization by an AM fungus could have an impact on root-associated microbial communities, being these variations potentially contributing to improve plant performance. Thereto, we determined the root-associated microbiomes in mycorrhizal rice roots and compared it to non-mycorrhizal plants. The AM-inoculated rice plants were grown simultaneously in the same paddy field as non-inoculated plants (see **Additional file 1: Fig. S1** for details on field assays). At the Class level, the taxonomic composition of root-compartments of mycorrhizal and non-mycorrhizal plants was similar, but the rice genotype and root compartment had an effect on the bacterial and fungal community in mycorrhizal roots (**Additional file 1: Fig. S5 and Fig. S6**)

Analysis of compositions of microbiomes (ANCOM) [38] was used to identify differences between mycorrhizal and non-mycorrhizal roots in the community of bacteria residing in the endosphere and rhizosphere in each genotype at the species-level resolution. In particular, ANCOM-BC2 (ANCOM with Bias Correction) is considered a conservative method for the analysis of differential abundance in microbiome studies designed to effectively control the false discovery rate (FDR). The endosphere of mycorrhizal Bomba plants was enriched in *Kosakonia*, *Arcobacter*, *Acinetobacter*, *Shewanella* and *Dickeya* species (**Fig. 5A**, orange bars), whereas the endosphere of non-mycorrhizal Bomba plants was enriched in *Klebsiella pasteurii* and *Streptococcus koreensis* (**Fig. 5A**, blue bars). Regarding JSendra plants, *Arcobacter* and *Shewanella* species were the most differentially abundant species in the endosphere of mycorrhizal roots (**Fig. 5B**, orange bars), while *Pseudomonas*, *Citrobacter, Phytobacter or Enterobacter* species were enriched in non-mycorrhizal JSendra roots (**Fig. 5B**, blue bars).

**Fig. 5.**
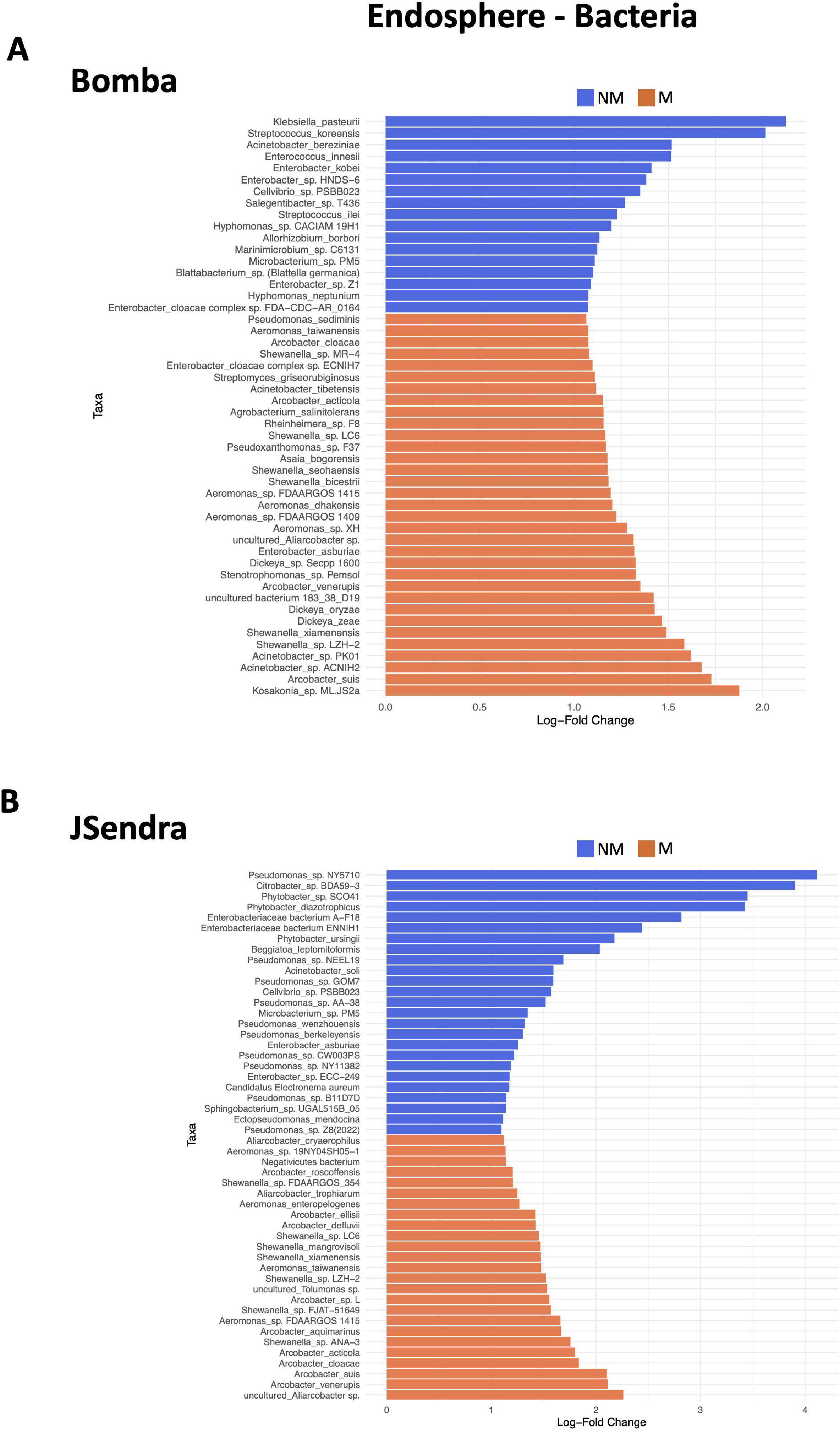
Specific bacterial signatures in mycorrhizal and non-mycorrhizal endosphere at the species level. ANCOM analysis was used to identify the 50 most differentially abundant bacterial species in the root endosphere of non-mycorrhizal and mycorrhizal Bomba (**A**) and JSendra (**B**) plants. Significantly enriched bacterial species at a p value < 0.05 (adjusted by Holm-Bonferroni method) are presented for each comparison. Blue, bacterial species enriched in the endosphere of non-mycorrhizal (NM) roots. Orange, bacterial species enriched in the endosphere of mycorrhizal (M) roots. Bars represent the relative abundance of each bacteria within the Bacteria Kingdom.

In the context of bacterial taxa contributing the most to the dissimilarity between mycorrhizal and non-mycorrhizal roots in the rhizosphere of rice varieties, *Klebsiella, Limnobacter*, *Trichormus* (as well as *Sulfurospirillum* and *Rheinheimera* species, among others), were found to preferentially colonize the rhizosphere of mycorrhizal Bomba plants (**Fig. 6A**, orange bars), whereas *Rickettsia*, *Planktothricoides*, *Streptococcus* and *Flavobacterium* were highly enriched in the rhizosphere of mycorrhizal JSendra plants (**Fig. 6B**, orange bars). Specific bacterial signatures in the rhizosphere of non-mycorrhizal roots included *Prevotella*, *Streptococcus* and *Rickettsia* species in Bomba plants (**Fig. 6A**, blue bars) and *Xanthomonas* and *Streptomyces* species in JSendra plants (**Fig. 6B**, blue bars).

**Fig. 6.**
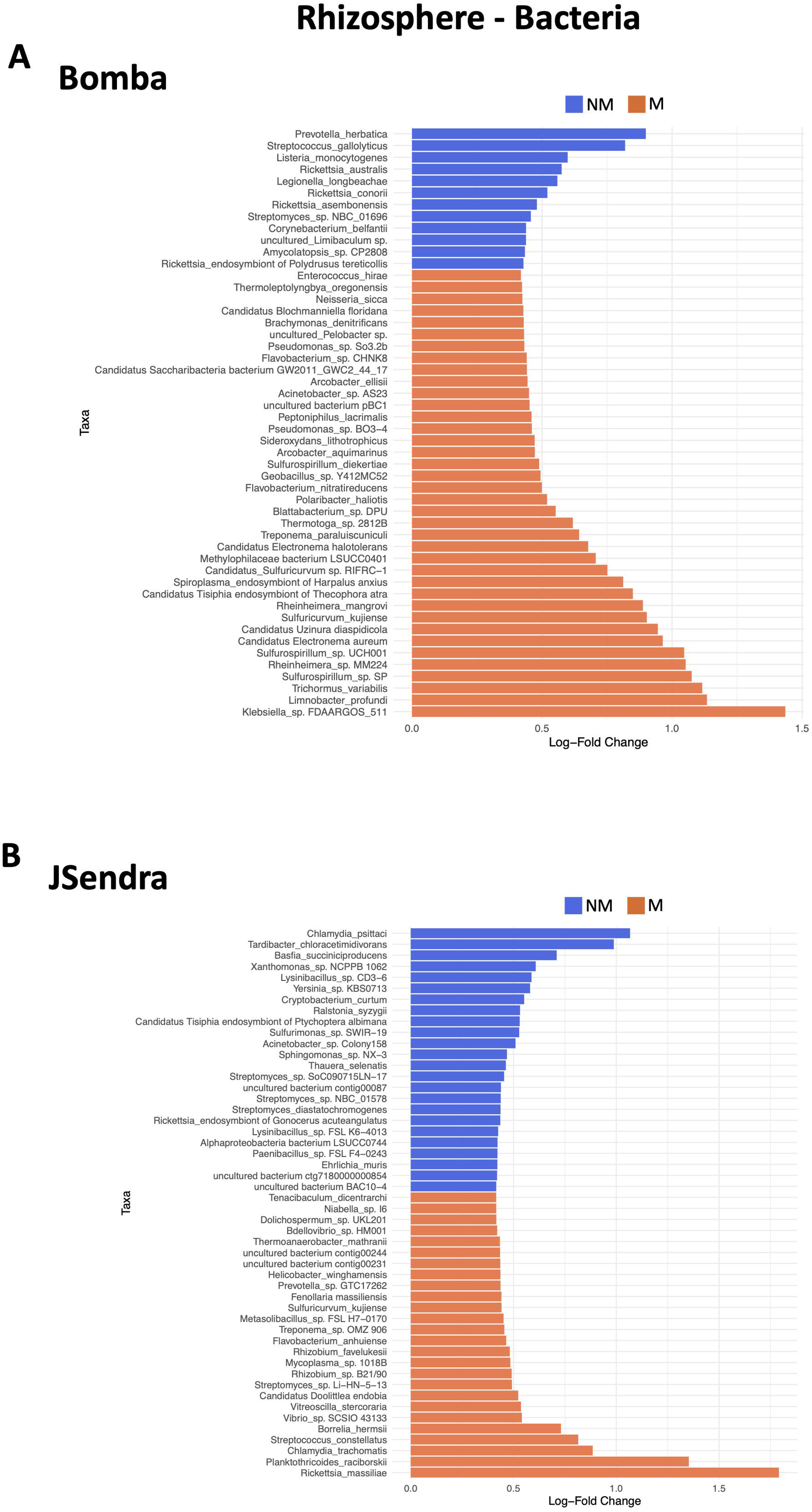
Specific bacterial signatures in mycorrhizal and non-mycorrhizal rhizosphere at the species level. ANCOM analysis was used to identify the 50 most differentially abundant bacterial species in the rhizosphere of non-mycorrhizal and mycorrhizal Bomba (**A**) and JSendra (**B**) plants. Significantly enriched bacterial species at a p value < 0.05 (adjusted by Holm-Bonferroni method) are presented for each comparison. Blue, bacterial species enriched in the rhizosphere of non-mycorrhizal (NM) roots. Orange, bacterial species enriched in the rhizosphere of mycorrhizal (M) roots. Bars represent the relative abundance of each bacteria within the Bacteria Kingdom.

The observation that *Xanthomonas* was identified in the rhizosphere of JSendra plants raised an interesting question. Among *Xanthomonas* species, *X. oryzae* causes Bacterial Leaf Blight (BLB), a devastating bacterial disease of rice worldwide. This bacteria lives in soil and enters into the roots through wounds. Once inside, the bacteria move upwards through the vascular system causing systemic infection into the aerial part of the plant. At present, however, no incidence of BLB has been reported in the rice-growing region in which rice plants were grown (Valencia, eastern region of the Iberian Peninsula). Close examination of *X. oryzae* reads in our metagenomic data sets revealed the presence of *X. oryzae* in the endosphere of Bomba and JSendra roots, its abundance being higher in the later one (**Additional file 1: Supplementary Figure 7**). Importantly, *X. oryzae* abundance decreased in mycorrhizal roots relative to non-mycorrhizal roots in both rice varieties. This observation suggest that the AM symbiosis might be beneficial for the host plant in preventing *X. oryzae* invasion of rice plants, and aspect that deserves further investigation.

Collectively, this analysis confirms that the arbuscular mycorrhizal symbiosis alters the bacterial root microbiome of rice plants grown under flooded conditions. This effect can be observed at both the endosphere and rhizosphere.

### Methanogens in root-associated microbiomes of rice plants

After exploring the composition of microbiomes, we put the focus on *bona fide* species related to functional activities. For instance, flooded rice fields are one of the major sources of biogenic methane which is mostly produced by methanogenic microorganisms in anaerobic conditions. To produce methane, methanogens utilize fertilizers, dead plant tissues, and organic exudates from plants. *Archaea* are the primary organisms that produce methane, including *Methanosarcina*, *Methanobacterium, Methanosaeta, Methanocella, Methanococcus* and *Methanospirillum*.

Our metagenomic data revealed the presence of different species of methanogenic archaea in the rhizosphere and root endosphere of rice plants (**Additional file 2: Table S4**). Species of the *Methanosarcina* genus, namely *M. siciliae* and *M. barkeri*, were the most abundant, methanogens in root compartments in both Bomba and JSendra plants, in both mycorrhizal and non-mycorrhizal roots (**Fig. 7A**). Of them, *M. siciliae* was more abundant than *M. barkeri* (**Fig. 7A**). In non-mycorrhizal plants, both the host genotype and the root compartment influenced the abundance of certain methanogenic species (**Fig. 7A**, left panel).

**Fig. 7.**
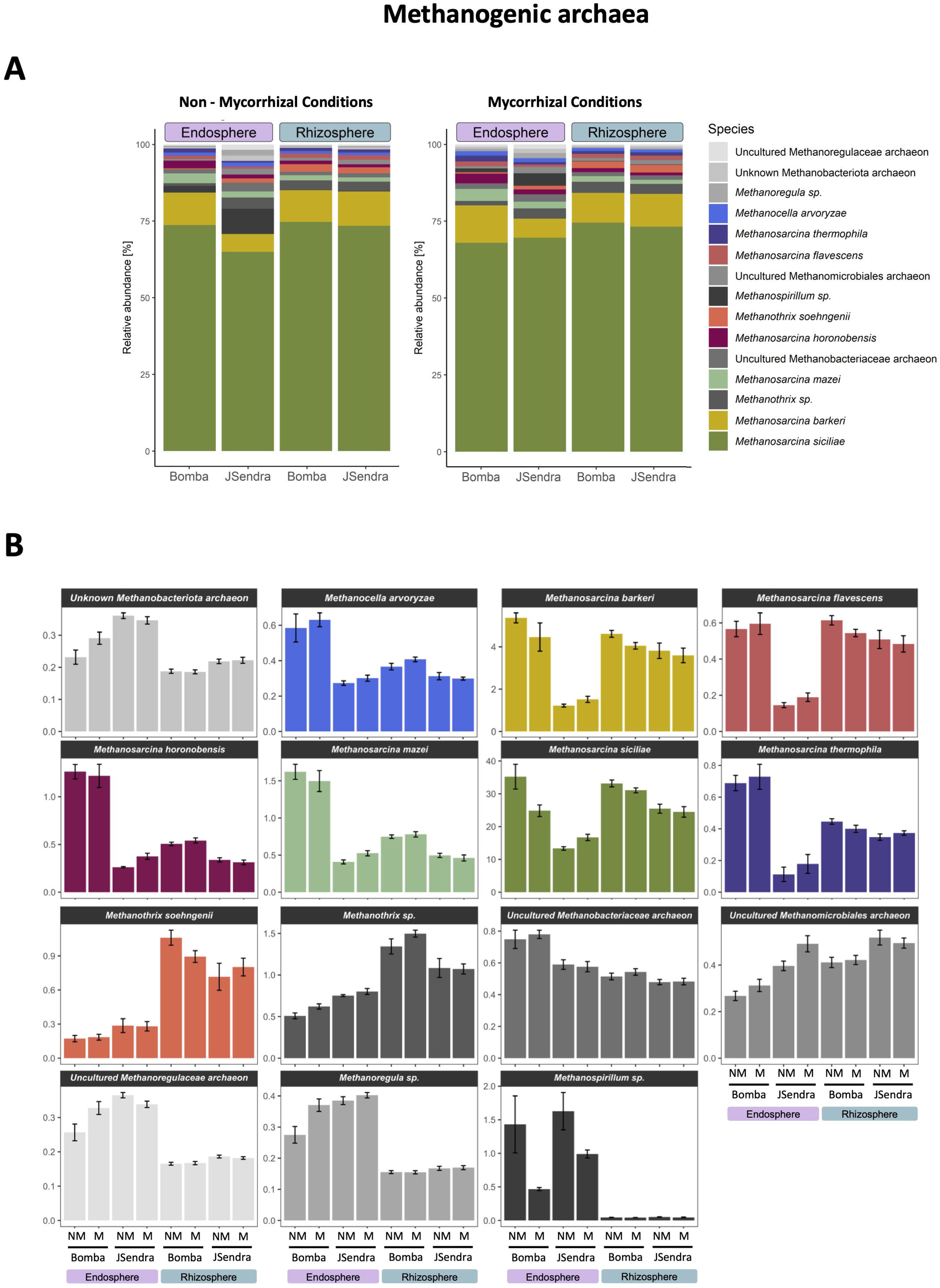
Relative abundance of Methanogenic archaea in the endosphere and rhizosphere of mycorrhizal and non-mycorrhizal rice plants. (**A**) Relative abundance within Methanogenic archaea at the species level in the root endosphere and soil rhizosphere of Bomba and JSendra plants (mycorrhizal and non-mycorrhizal condition in each variety). (**B**) Relative abundance of each Methanogenic archaea within the Archaea kingdom.

As shown in **Fig. 7B**, *Methanocella arvoryzae*, *Methanosarcina horonobensis* and *Methanosarcina thermophila* were identified predominantly in the endosphere of Bomba plants. *M. barkeri* and *Methanosarcina flavescens* were less abundant in the endosphere of JSendra plants (**Fig. 7B**). Considering root compartments, *Methanothrix soehngenii* was present mostly in the rhizosphere in both varieties (**Fig. 7B**). Finally, the mycorrhizal status of the rice plant also had an effect on the abundance of certain methanogenic Archaea species. In both rice varieties, *Methanospirillum sp* was more abundant in the endosphere of non-mycorrhizal roots than in the endosphere of mycorrhizal roots (**Fig. 7B**).

Shotgun metagenomics also identified a series of uncultured methanogenic archaea belonging to different taxa (e.g., uncultured *Methanoregulaceae archaeon*) whose abundance was higher in the root endosphere in both rice varieties (non-mycorrhizal and mycorrhizal conditions) (**Fig. 7B**).

Together, these results suggest that in regard to methanogens, the host genotype and root compartment play again a key role to determine the structure of the root microbiome. However, the association of rice roots with the AM fungus could alter the balance of certain species.

### Nutrient cycling bacteria in root-associated microbiomes of rice varieties

Another central functional role of microbial communities in aquatic ecosystems is nutrient cycling by facilitating the conversion of organic matter into accessible forms of essential elements. Importantly, our understanding on bacteria involved in nutrient cycling in paddy rice fields is still limited. Regarding P cycling, it should be mentioned that in agricultural soils only a small fraction of P is available for assimilation by plants due to immobilization of this element in different complexes, and an important fraction of Pi applied in fertilizers cannot be assimilated by plants [39]. However, certain microorganisms can make P bioavailable to plants by hydrolyzing organic and inorganic P compounds into soluble forms, thus making P bioavailable to plants. Phosphate solubilizing microorganisms include bacteria belonging to different genera, generally known as Phosphate Solubilizing Bacteria (PSB) [39, 40]. Reported PSB include genera such as *Bacillus*, *Pseudomonas*, *Rhizobium*, *Enterobacter*, *Burkholderia*, *Achromobacter, Micrococcus*, *Flavobacterium*, and *Erwinia*, with *Pseudomonas*, *Bacillus* and *Rhizobium* being the most powerful phosphate solubilizers [41–44]. On the other hand, bacterial species are also considered to be major drivers of N-cycling processes (herein after referred to as NCyc bacteria) in flooded rice fields. Nitrification-denitrification processes are dependent on the activities of different bacteria, including species from the *Rhizobium* genus which promote nitrogen fixation by converting atmospheric nitrogen into ammonia.

The heatmap in **Fig. 8** summaries the differential abundance of PSB and NCyc bacteria of rice plants in both mycorrhizal and non-mycorrhizal plants grown in paddy fields. Details on the relative abundance of PSB and NCyc in the different samples are presented in **Additional file 2: Table S5**. Briefly, the two first clusters depict rhizosphere-(cluster I) and endosphere-specific bacteria (cluster II) that showed almost identical abundance in both genotypes. Note that the rhizosphere cluster encompassed specialised PSB and NCyc bacteria, while the endosphere cluster included bacteria capable to operate as both PSB and NCyc. The rest of the clusters consisted of bacteria that were more prominent within endosphere depending on the genotype (JSendra, cluster III; Bomba, cluster IV) and also could work as PSB and NCyc. Two thirds of the total bacteria were allocated at the endosphere, highlighting the relevance of the diversifying bacteria specialised in nutrient cycling withing the root. Remarkably, AM-colonisation led to lowered abundance for the genotype-specific species in clusters III and IV – a fact previously observed for methanogen species. These patterns allowed us to propose a selection for endosphere-resident bacteria based on versatility to cycle for both P and N, whereas rhizosphere bacteria could be more specialised for one or another mineral. Moreover, they reinforce the scenario whereby AM-root colonisation led to diminish bacteria contributing to plant nutrition.

**Fig. 8.**
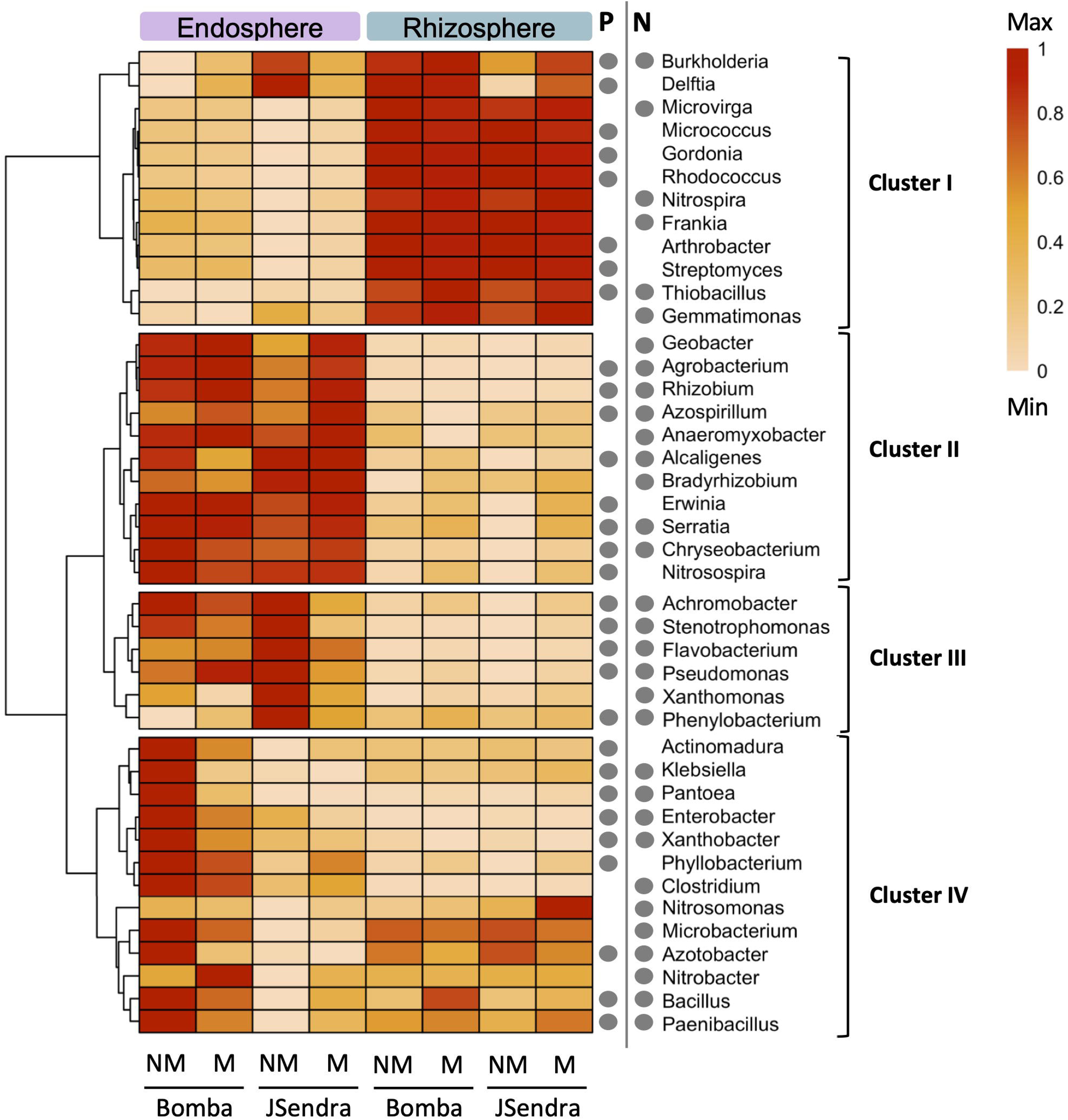
PSB and NCyc genera found in the endosphere and rhizosphere of rice plants. (**A**) Heatmap showing the enrichment of PSB and NCyc genera in both Bomba and JSendra plants. (**B**) Relative abundance of selected PSB and NCyc bacteria within the Bacteria Kingdom. NM, non-mycorrhizal plants; M, mycorrhizal plants; P, Phosphate; PSB, Phosphate Solubilizing Bacteria; N, Nitrogen; NCyc, Nitrogen Cycling bacteria.

## Discussion

The present study provides a comprehensive characterization of microbiomes in root-associated compartments in rice plants growing in flooded fields by means of shotgun metagenomic sequencing. Importantly, we put the focus on two temperate *japonica* rice varieties of agronomical interest, Bomba and JSendra, which are well adapted to Mediterranean climate conditions. Metagenomic analyses were conducted in roots of adult rice plants supporting that microbial communities identified in this study reflect microbial taxa that have been shaped along the plant’s life cycle through selection of specialized microbial taxa.

Our analysis revealed that the root compartment is the main factor that contributes to shape the microbial communities in rice roots. Quantitatively, bacterial taxa showed a much greater abundance and heterogeneity than fungal taxa across genotypes and root compartments (**Fig. 1**). In both genotypes, the bacterial microbiomes were particularly enriched in Pseudomonadota and Actinomycetota. In the Phylum Pseudomonadota, the prevalent Classes were Alpha*-*, Beta- and Gamma*-*Proteobacteria, their relative abundance being notably higher in the endosphere than in the rhizosphere compartment in both varieties. Conversely, Actinomycetes were more abundant in the rhizosphere than in the endosphere. Regarding the fungal community, *Ascomycota* and *Basidiomycota* dominated root-associated compartments in rice plants. Within the Phylum *Ascomycota*, *Sordariomycetes* were more abundant in the endosphere than in the rhizosphere, while *Eurotiomycetes* were more prevalent in rhizosphere. The differences between compartments were even more remarkable for bacterial classes in minority. The preferential colonization of different compartments by distinct bacterial and fungal species supports the capacity of rice plants to actively recruit the root microbiome from the rhizosphere. Moreover, some components of the main communities in endosphere have been previously reported as beneficial species. For instance, within *Sordariomycetes*, *Chaetomium* species are considered to be promising biological control agents against rice diseases with important applications in rice straw degradation [45]. The N-fixing *Rhizobium*, a symbiotic bacteria capable of invading roots on leguminous plants to improve nutrient uptake and growth, also occur as endophytes in rice roots [46, 47]. *Flavobacterium anhuiense* contributes to potassium solubilizing [48] and *Pseudomonas sediminis* and *Agrobacterium salinitolerans* are salt tolerant bacteria related to nutrient mobilisation and inhibiting the growth of certain pathogenic fungi [49, 50]. Towards understanding how plants could mechanistically select for beneficial microbiomes, increasing evidence support that variations in rhizosphere microbiomes are due to differences in the composition of root exudates in aerobic soil [51], though additional factors cannot be ruled out.

In a second term, we provide evidence that the host genotype can further determine the composition of the microbiomes. In our study, microbial community variations between Bomba and JSendra plants were more evident at the species level.

We would like to remark that our results partially differ from previous reports on microbial communities associated with rice roots [6, 10–12]. We attribute the discrepancies to the biogeography and cultivation systems of rice varieties, environmental factors, and/or host genotypes and the fact that microbial taxa identified in Bomba and JSendra roots represent well-adapted species to these rice varieties that can establish associations with rice plants in paddy fields, at least in the western Mediterranean region where both varieties are widely grown. In consequence, our study evidences the urgency to conduct further characterisation of microbiomes for local rice varieties in the context of the corresponding cultivar areas and growth conditions.

### Differences in bacterial assemblages associated with methane production and nutrient solubilization in roots of rice plants under flooded conditions

Beyond identifying differentially abundant taxonomic groups in root-associated compartments of rice plants, we wanted to address whether the microbiome of rice roots could contribute to adaptation to the conditions where rice is cultivated, that is, flooded conditions. This motivation was further reinforced by the autonomous potential of rice plants to shape the composition of the microbiome we found. Thereto, our metagenomic dataset was dissected for *bona fide* microorganisms related to methane production and nutrient recycling.

In flooded rice fields, methane is produced by methanogenic microbes. Methanogenic Archaea *Methanosarcina* emerged as the prevalent genus in the endosphere and rhizosphere of both Bomba and JSendra plants. Although the relative abundance of species at the rhizosphere tended to remain steady, the composition at the endosphere was shaped by the genotype. Thus, the endosphere of Bomba roots had increased abundance in *M*. *horonobensis, M. thermophila, M. flavescens* and *M. barkeri* compared to JSendra. On the other hand, JSendra root endosphere contained more *Methanoregulaceae archeon* and *Methanobacteriota archeon* species. This finding further supported by the ability of the host to select their microbiome, prompted us to speculate that a differential composition of methanogenic microbes in Bomba and JSendra might have an impact on methane production. Clearly, a reduction in methane emissions from paddy fields is desirable, hence, knowing the population of methanogenic microbes in rice varieties grown in paddy fields will help in this direction.

Our analyses also showed a common compartment-specific signature for the bacterial community involved in nutrient cycling, particularly PSB and NCyc bacteria, between the two genotypes (**Fig. 8**). To note, common rhizosphere communities were more specialised to either PSBs or NCyc, while those in the endosphere in both rice varieties could mainly participate in P and N cycling. This last pattern could be also observed for other genotype-specific endosphere bacteria. Accordingly, we propose an economization strategy whereby host plants have developed mechanisms to prioritize associations with bacteria with higher versatility to mobilise nutrients to ultimately maximise the benefits of the colonisation.

### Variations in the root microbiome in mycorrhizal rice plants in flooded conditions

Root colonization by AM fungi in aerobic rice has long been demonstrated [52, 53]. In paddy fields, however, flooding appears to inhibit root colonization by AM fungi [54]. Importantly, we previously reported pre-inoculation with *R. irregularis* and early growth of rice plants under aerobic conditions prior transplanting to paddy fields allows an effective symbiotic relationship in flooded rice, thus, supporting that once the AM fungus has penetrated into the root system, the functionality of the AM symbiosis is not affected by flooding [18, 19]. Most importantly, root colonization by *R. irregularis* increases grain yield and enhances blast resistance in rice plants in paddy fields, also in Bomba and JSendra plants [19]. Samples used for metagenomic analyses in this study were harvested from Bomba and JSendra plants from field assays described by [19]. To complement our study, the impact of AM fungi colonization in the root microbiome was evaluated.

Results here presented revealed differences in genotype-specific microbiome of mycorrhizal roots relative to non-mycorrhizal roots of rice plants, with an overall trend to reduce their abundance. In particular, the lowering of genotype-specific bacteria involved in phosphate solubilization and nutrient cycling PSB bacteria would be in line with the necessity of avoid high Pi availability to sustain AM-colonisation. Earlier studies reported a cooperative relationship between AM fungi and PSB in soil making P more available to the plant and/or increasing tolerance to stress [55–58]. Given the beneficial consequences of AM colonization we contemplate three plausible scenarios: (i) AM fungi could efficiently compete with the other potential colonisers; (ii) the homeostatic state upon AM colonisation might alter or interfere with the plant signalling mechanisms to attract microbiota; or (iii) the host might actively focus the recruiting strategy on AM fungi, as the most advantageous case.

A priori, the root microbial composition reflects the outcome of complex interactions involving the host plant and microbe-microbe interactions, and root colonization by an AM fungus might influence root colonization by distinct microbes. Microbial assemblages can then be influenced by the mycorrhizal status of the rice plant, as illustrated in this study for *X. oryzae* in the endosphere of Bomba and JSendra plants. Through the comparison of abundances in microbial species in non-mycorrhizal and mycorrhizal roots, it was possible to discern that the abundance of *X. oryzae* decreased in mycorrhizal roots relative to non-mycorrhizal roots. Microbial species whose abundance was modulated in mycorrhizal roots included species that can be beneficial to the host plant, such as *Rhizobium* (enriched in mycorrhizal roots of JSendra relative to non-mycorrhizal roots) supporting that specific bacterial species are selected in mycorrhizal rice roots.

## Conclusion

This study provides evidence on genotype-specific and root compartment-specific differences in the microbial communities in rice plants growing under flooded conditions. Not only the host genotype and root compartment, but also the mycorrhizal condition can shape the rice root microbiome. Bacteria constitute the vast majority of microbial taxa in the rice root, making up approximately 95% of the total microbia. The greatest variations in both bacterial and fungal taxa were observed at species level. Nowadays is generally accepted that microbial assemblages exert a great influence on plant growth and productivity while contributing to adaptation to environmental challenges. Although the application of microbial consortia has proven to be useful in stimulating plant growth and alleviating environmental stresses in different plant species, in rice (particularly in flooded rice), we are only at the beginning of understanding the complexity of plant-microbe and microbe-microbe associations. The information here presented on the root-associated microbiomes offers practical guidance for the development of sustainable strategies for microbiome-assisted rice cultivation while reducing reliance on agrochemical fertilizers in paddy fields.

## Declarations

### Ethics approval and consent to participate

Not applicable.

### Consent for publication

Not applicable.

### Availability of data and materials

The original data presented in this study are publicly available in the European Nucleotide Archive (ENA, http://www.ebi.ac.uk/ena) at the EMBL-EBI under accession PRJEB105165.

### Author contributions

BSS designed research; IB and HM-C prepared and analysed plant materials; CD, contributed to field trials and sample harvesting; IB, HM-C and AG-M formal analysis of metagenomic data and interpretation of results; BSS, Writing original draft and funding acquisition; All authors, review and editing. All authors read and approved the final manuscript.

## Acknowledgements

We thank members of the laboratory Laia Castillo and Gerrit Bücker for assistance in sample preparation. We also thank Victor M. González from CRAG’s bioinformatic unit for technical assistance in metagenomic analysis. Héctor Martín-Cardoso was a recipient of a grant from the Ministerio de Ciencia e Innovación (PRE2019–087477).

## Supplementary Information

**Additional file 1: Fig. S1. Experimental design followed to grow rice plants in flooded rice fields**. Distribution of plots containing the rice varieties Bomba and JSendra, mock-inoculated and *R. irregularis* inoculated.

**Additional file 1: Fig. S2. Proportion of reads assigned to Fungi, Bacteria, Archaea and Viruses in root endosphere and rhizosphere**. Bacteria, 94-96%, Fungi, 2-3%, Archaea, 1-2%, Viruses, 0.3%.

**Additional file 1: Fig. S3. Differential abundance of microbial taxa in the root endosphere and rhizosphere of non-mycorrhizal Bomba and JSendra plants**. Data is presented at the species level. Heatmap represents bacterial and fungal species with a relative abundance greater than 0.01% in each sample. The Min-Max scaling normalization method was applied to the data after performing the logD (xD+D1) transformation, rescaling it to the 0 to 1 range. The colour code indicates differences in the relative abundance ranging from light (less abundant) to dark (more abundant) red.

**Additional file 1: Fig. S4. The most abundant fungal taxa identified in the rhizosphere of Bomba (A) and JSendra (B) plants**. Sankey diagrams showing the relative abundance of the top 20 taxa in each variety is presented, from the kingdom (on the left) to species (on the right). The width of the bar is proportional to the abundance, whereas nodes depict the hierarchy levels of microbial taxa.

**Additional file 1: Fig. S5. Taxonomic composition of the bacterial and fungal communities in mycorrhizal roots of Bomba and JSendra plants**. Different colors indicate different taxa at the Class level.

**Additional file 1: Fig. S6. Differential abundance of bacterial and fungal taxa in the root endosphere and rhizosphere of mycorrhizal Bomba and JSendra plants**. Different colors indicate different taxa at the species level. Heatmap represents species with a relative abundance greater than 0.01% in each sample. The Min-Max scaling normalization method was applied to the data after performing the logD (xD+D1) transformation, rescaling it to the 0 to 1 range. The colour code indicates differences in the relative abundance ranging from light (less abundant) to dark (more abundant) red.

**Additional file 1: Fig. S7. Relative abundance of Xanthomonas oryzae.** Data for non-mycorrhized and mycorrhized conditions from Bomba and JSendra is shown.

**Additional file 2: Table S1. Statistics for data pre-processed data**. Each sample corresponds to the average of 3 technical replicates.

**Additional file 2: Table S2. Complete list of species identified in metagenomic data from the root endosphere and rhizosphere.** Data show the absolute abundance of every species identified in each compartment, rice genotype (Bomba, JSendra), and condition (M, mycorrhizal; NM, non-mycorrhizal).

**Additional file 2: Table S3. Relative abundance of bacterial and fungal taxa in the root endosphere and rhizosphere of Bomba and JSendra plants**. Data indicate the percentage of each Class in all Bacteria.

**Additional file 2: Table S4**. **List of methanogenic archaea identified in the endosphere and rhizosphere of Bomba and JSendra plants**. The relative abundance of each species in the Phylum Methanobacteriota, Thermoplasmatota and Thermoproteota among Methanogenic Archaea) is shown.

**Additional file 2: Table S5**. **List of Phosphate Solubilizing Bacteria (PSB) and Nitrogen Cycling (NCyc) bacteria species identified in the endosphere and rhizosphere of Bomba and JSendra plants**. Data show relative abundance among bacteria.

## References

1. Aminurrasyid AH Bin, Mohd Ikmal A, Nadarajah KK. The Rice-Microbe Nexus: Unlocking Productivity Through Soil Science. Rice (N Y). 2025;18:56. 10.1186/s12284-025-00809-0.

2. He X, Batáry P, Zou Y, Zhou W, Wang G, Liu Z, et al. Agricultural diversification promotes sustainable and resilient global rice production. Nat Food. 2023;4:788–96. 10.1038/s43016-023-00836-4.

3. Kim H, Lee Y-H. The Rice Microbiome: A Model Platform for Crop Holobiome. Phytobiomes J. 2020;4:5–18. 10.1094/PBIOMES-07-19-0035-RVW.

4. Compant S, Cassan F, Kostić T, Johnson L, Brader G, Trognitz F, et al. Harnessing the plant microbiome for sustainable crop production. Nat Rev Microbiol. 2025;23:9–23. 10.1038/s41579-024-01079-1.

5. Misu IJ, Kayess MdO, Siddiqui MdN, Gupta DR, Islam MN, Islam T. Microbiome Engineering for Sustainable Rice Production: Strategies for Biofertilization, Stress Tolerance, and Climate Resilience. Microorganisms. 2025;13:233. 10.3390/microorganisms13020233.

6. Ding L-J, Cui H-L, Nie S-A, Long X-E, Duan G-L, Zhu Y-G. Microbiomes inhabiting rice roots and rhizosphere. FEMS Microbiol Ecol. 2019;95:fiz040. 10.1093/femsec/fiz040.

7. Luo X, Fu X, Yang Y, Cai P, Peng S, Chen W, et al. Microbial communities play important roles in modulating paddy soil fertility. Sci Rep. 2016;6:20326. 10.1038/srep20326.

8. Ramírez-Viga TK, Aguilar R, Castillo-Argüero S, Chiappa-Carrara X, Guadarrama P, Ramos-Zapata J. Wetland plant species improve performance when inoculated with arbuscular mycorrhizal fungi: a meta-analysis of experimental pot studies. Mycorrhiza. 2018;28:477–93. 10.1007/s00572-018-0839-7.

9. Kumar M, Kumar A, Sahu KP, Patel A, Reddy B, Sheoran N, et al. Deciphering core-microbiome of rice leaf endosphere: Revelation by metagenomic and microbiological analysis of aromatic and non-aromatic genotypes grown in three geographical zones. Microbiol Res. 2021;246:126704. 10.1016/j.micres.2021.126704.

10. Dastogeer KMG, Yasuda M, Okazaki S. Microbiome and pathobiome analyses reveal changes in community structure by foliar pathogen infection in rice. Front Microbiol. 2022;13:949152. 10.3389/fmicb.2022.949152.

11. Zhang X, Ma Y-N, Wang X, Liao K, He S, Zhao X, et al. Dynamics of rice microbiomes reveal core vertically transmitted seed endophytes. Microbiome. 2022;10:216. 10.1186/s40168-022-01422-9.

12. Juliyanti V, Itakura R, Kotani K, Lim SY, Suzuki G, Chong CW, et al. Comparative analysis of root associated microbes in tropical cultivated and weedy rice (Oryza spp.) and temperate cultivated rice. Sci Rep. 2024;14:9656. 10.1038/s41598-024-60384-0.

13. Islam MM, Jana SK, Sengupta S, Mandal S. Impact of Rhizospheric Microbiome on Rice Cultivation. Curr Microbiol. 2024;81:188. 10.1007/s00284-024-03703-y.

14. Tan LT, Dailin DJ, Hanapi SZ, Rahman RA, Mehnaz S, Shahid I, et al. Role of Microbiome on Healthy Growth and Yield of Rice Plant. 2024. p. 141–61. 10.1007/978-981-99-9388-8_9.

15. Dastogeer KMG, Tumpa FH, Sultana A, Akter MA, Chakraborty A. Plant microbiome–an account of the factors that shape community composition and diversity. Curr Plant Biol. 2020;23:100161. 10.1016/j.cpb.2020.100161.

16. Edwards J, Johnson C, Santos-Medellín C, Lurie E, Podishetty NK, Bhatnagar S, et al. Structure, variation, and assembly of the root-associated microbiomes of rice. Proc Natl Acad Sci USA. 2015;112:E911–20. 10.1073/pnas.1414592112.

17. Moronta-Barrios F, Gionechetti F, Pallavicini A, Marys E, Venturi V. Bacterial Microbiota of Rice Roots: 16S-Based Taxonomic Profiling of Endophytic and Rhizospheric Diversity, Endophytes Isolation and Simplified Endophytic Community. Microorganisms. 2018;6:14. 10.3390/microorganisms6010014.

18. Campo S, Martín-Cardoso H, Olivé M, Pla E, Catala-Forner M, Martínez-Eixarch M, et al. Effect of Root Colonization by Arbuscular Mycorrhizal Fungi on Growth, Productivity and Blast Resistance in Rice. Rice. 2020;13:42. 10.1186/s12284-020-00402-7.

19. Martín-Cardoso H, Castillo L, Busturia I, Bücker G, Marqués L, Pla E, et al. Arbuscular Mycorrhizal Fungi Increase Blast Resistance and Grain Yield in Japonica Rice Cultivars in Flooded Fields. Rice. 2025;18:47. 10.1186/s12284-025-00805-4.

20. Murray MG, Thompson WF. Rapid isolation of high molecular weight plant DNA. Nucleic Acids Res. 1980;8:4321–6. 10.1093/nar/8.19.4321.

21. Andrews S. FastQC: A Quality Control Tool for High Throughput Sequence Data. 2010.

22. Chen S, Zhou Y, Chen Y, Gu J. fastp: an ultra-fast all-in-one FASTQ preprocessor. Bioinformatics. 2018;34:i884–90. 10.1093/bioinformatics/bty560.

23. Langmead B, Wilks C, Antonescu V, Charles R. Scaling read aligners to hundreds of threads on general-purpose processors. Bioinformatics. 2019;35:421–32. 10.1093/bioinformatics/bty648.

24. Wood DE, Lu J, Langmead B. Improved metagenomic analysis with Kraken 2. Genome Biol. 2019;20:257. 10.1186/s13059-019-1891-0.

25. Lu J, Breitwieser FP, Thielen P, Salzberg SL. Bracken: estimating species abundance in metagenomics data. PeerJ Comput Sci. 2017;3:e104. 10.7717/peerj-cs.104.

26. Lahti L, Shetty S. microbiome R package. Bioconductor. 10.18129/B9.bioc.microbiome.

27. McMurdie PJ, Holmes S. phyloseq: An R Package for Reproducible Interactive Analysis and Graphics of Microbiome Census Data. PLoS One. 2013;8:e61217. 10.1371/journal.pone.0061217.

28. Wickham H, Averick M, Bryan J, Chang W, McGowan L, François R, et al. Welcome to the Tidyverse. J Open Source Softw. 2019;4:1686. 10.21105/joss.01686.

29. Martinez-Arbizu P. pairwiseAdonis: Pairwise multilevel comparison using adonis. 2020.

30. Oksanen J, Simpson GL, Blanchet FG, Kindt R, Legendre P, Minchin PR, et al. vegan: Community Ecology Package. CRAN: Contributed Packages. 2001. 10.32614/CRAN.package.vegan.

31. Shannon CE, Weaver W. The Mathematical Theory of Communication. Urbana, IL: The University of Illinois Press; 1964.

32. Margalef R. Information Theory in Ecology. General Systems. 1958;3:36–71.

33. Han J, Kamber M, Pei J. Data mining: concepts and techniques. 3rd Edition. Morgan Kaufmann; 2011.

34. Breitwieser FP, Salzberg SL. Pavian: interactive analysis of metagenomics data for microbiome studies and pathogen identification. Bioinformatics. 2020;36:1303–4. 10.1093/bioinformatics/btz715.

35. Lin H, Peddada S Das. Multigroup analysis of compositions of microbiomes with covariate adjustments and repeated measures. Nat Methods. 2024;21:83–91. 10.1038/s41592-023-02092-7.

36. Gan HM, Lee YP, Austin CM. Nanopore Long-Read Guided Complete Genome Assembly of Hydrogenophaga intermedia, and Genomic Insights into 4-Aminobenzenesulfonate, p-Aminobenzoic Acid and Hydrogen Metabolism in the Genus Hydrogenophaga. Front Microbiol. 2017;8:1880. 10.3389/fmicb.2017.01880.

37. Liu Y, Tie B, Li Y, Lei M, Wei X, Liu X, et al. Inoculation of soil with cadmium-resistant bacterium Delftia sp. B9 reduces cadmium accumulation in rice (Oryza sativa L.) grains. Ecotoxicol Environ Saf. 2018;163:223–9. 10.1016/j.ecoenv.2018.07.081.

38. Mandal S, Van Treuren W, White RA, Eggesbø M, Knight R, Peddada SD. Analysis of composition of microbiomes: a novel method for studying microbial composition. Microb Ecol Health Dis. 2015;26:27663. 10.3402/mehd.v26.27663.

39. Silva LI da, Pereira MC, Carvalho AMX de, Buttrós VH, Pasqual M, Dória J. Phosphorus-Solubilizing Microorganisms: A Key to Sustainable Agriculture. Agriculture. 2023;13:462. 10.3390/agriculture13020462.

40. Zhu Y, Xing Y, Li Y, Jia J, Ying Y, Shi W. The Role of Phosphate-Solubilizing Microbial Interactions in Phosphorus Activation and Utilization in Plant–Soil Systems: A Review. Plants. 2024;13:2686. 10.3390/plants13192686.

41. Etesami H, Jeong BR, Glick BR. Contribution of Arbuscular Mycorrhizal Fungi, Phosphate–Solubilizing Bacteria, and Silicon to P Uptake by Plant. Front Plant Sci. 2021;12:699618. 10.3389/fpls.2021.699618.

42. Rawat P, Das S, Shankhdhar D, Shankhdhar SC. Phosphate-Solubilizing Microorganisms: Mechanism and Their Role in Phosphate Solubilization and Uptake. J Soil Sci Plant Nutr. 2021;21:49–68. 10.1007/s42729-020-00342-7.

43. Devi R, Kaur T, Kour D, Yadav A, Yadav AN, Suman A, et al. Minerals solubilizing and mobilizing microbiomes: A sustainable approach for managing minerals’ deficiency in agricultural soil. J Appl Microbiol. 2022;133:1245–72. 10.1111/jam.15627.

44. Pan L, Cai B. Phosphate-Solubilizing Bacteria: Advances in Their Physiology, Molecular Mechanisms and Microbial Community Effects. Microorganisms. 2023;11:2904. 10.3390/microorganisms11122904.

45. Song JJ, Soytong K, Kanokmedhakul S, Kanokmedhakul K, Poeaim S. Antifungal activity of microbial nanoparticles derived from Chaetomium spp. against Magnaporthe oryzae causing rice blast. Plant Protection Science. 2020;56:180–90. 10.17221/41/2019-PPS.

46. Biswas JC, Ladha JK, Dazzo FB. Rhizobia Inoculation Improves Nutrient Uptake and Growth of Lowland Rice. Soil Sci Soc Am J. 2000;64:1644–50. 10.2136/sssaj2000.6451644x.

47. Biswas JC, Ladha JK, Dazzo FB, Yanni YG, Rolfe BG. Rhizobial Inoculation Influences Seedling Vigor and Yield of Rice. Agron J. 2000;92:880–6. 10.2134/agronj2000.925880x.

48. Meena VS, Maurya BR, Verma JP, Aeron A, Kumar A, Kim K, et al. Potassium solubilizing rhizobacteria (KSR): Isolation, identification, and K-release dynamics from waste mica. Ecol Eng. 2015;81:340–7. 10.1016/j.ecoleng.2015.04.065.

49. Hao H, Wang Z, Gou C, Sha S, Yan C, Niu D, et al. Genome Sequence of the Agrobacterium salinitolerans DG3-1 Isolated from Cotton Roots. Mol Plant Microbe Interact. 2021;34:1458–60. 10.1094/MPMI-06-21-0154-A.

50. Liu Y, Yin P, Zhou J, Ma Y, Lai X, Lin J, et al. Removal of Nitrogen and Phosphorus by a Novel Salt-Tolerant Strain Pseudomonas sediminis D4. Water (Basel). 2025;17:502. 10.3390/w17040502.

51. Wankhade A, Wilkinson E, Britt DW, Kaundal A. A Review of Plant–Microbe Interactions in the Rhizosphere and the Role of Root Exudates in Microbiome Engineering. Applied Sciences. 2025;15:7127. 10.3390/app15137127.

52. Vallino M, Greppi D, Novero M, Bonfante P, Lupotto E. Rice root colonisation by mycorrhizal and endophytic fungi in aerobic soil. Ann appl Biol. 2009;154:195–204. 10.1111/j.1744-7348.2008.00286.x.

53. Bernaola L, Cange G, Way MO, Gore J, Hardke J, Stout M. Natural Colonization of Rice by Arbuscular Mycorrhizal Fungi in Different Production Areas. Rice Sci. 2018;25:169–74. 10.1016/j.rsci.2018.02.006.

54. Vallino M, Fiorilli V, Bonfante P. Rice flooding negatively impacts root branching and arbuscular mycorrhizal colonization, but not fungal viability. Plant Cell Environ. 2014;37:557–72. 10.1111/pce.12177.

55. Norouzinia F, Ansari MH, Aminpanah H, Firozi S. Alleviation of Soil Salinity on Physiological and Agronomic Traits of Rice Cultivars Using Arbuscular Mycorrhizal Fungi and Pseudomonas Strains Under Field Conditions. Rev Agric Neotrop. 2020;7:25–42. 10.32404/rean.v7i1.4042.

56. Nacoon S, Jogloy S, Riddech N, Mongkolthanaruk W, Ekprasert J, Cooper J, et al. Combination of arbuscular mycorrhizal fungi and phosphate solubilizing bacteria on growth and production of Helianthus tuberosus under field condition. Sci Rep. 2021;11:6501. 10.1038/s41598-021-86042-3.

57. Bao X, Zou J, Zhang B, Wu L, Yang T, Huang Q. Arbuscular Mycorrhizal Fungi and Microbes Interaction in Rice Mycorrhizosphere. Agronomy. 2022;12:1277. 10.3390/agronomy12061277.

58. Elekhtyar NM, Awad-Allah MMA, Alshallash KS, Alatawi A, Alshegaihi RM, Alsalmi RA. Impact of Arbuscular Mycorrhizal Fungi, Phosphate Solubilizing Bacteria and Selected Chemical Phosphorus Fertilizers on Growth and Productivity of Rice. Agriculture. 2022;12:1596. 10.3390/agriculture12101596.

