## Supplementary Figures for "Decoding microbial diversity in roots of rice plants under flooded conditions: influence of the host genotype, root compartment and mycorrhizal association"

### Additional file 1: Supplementary Figures

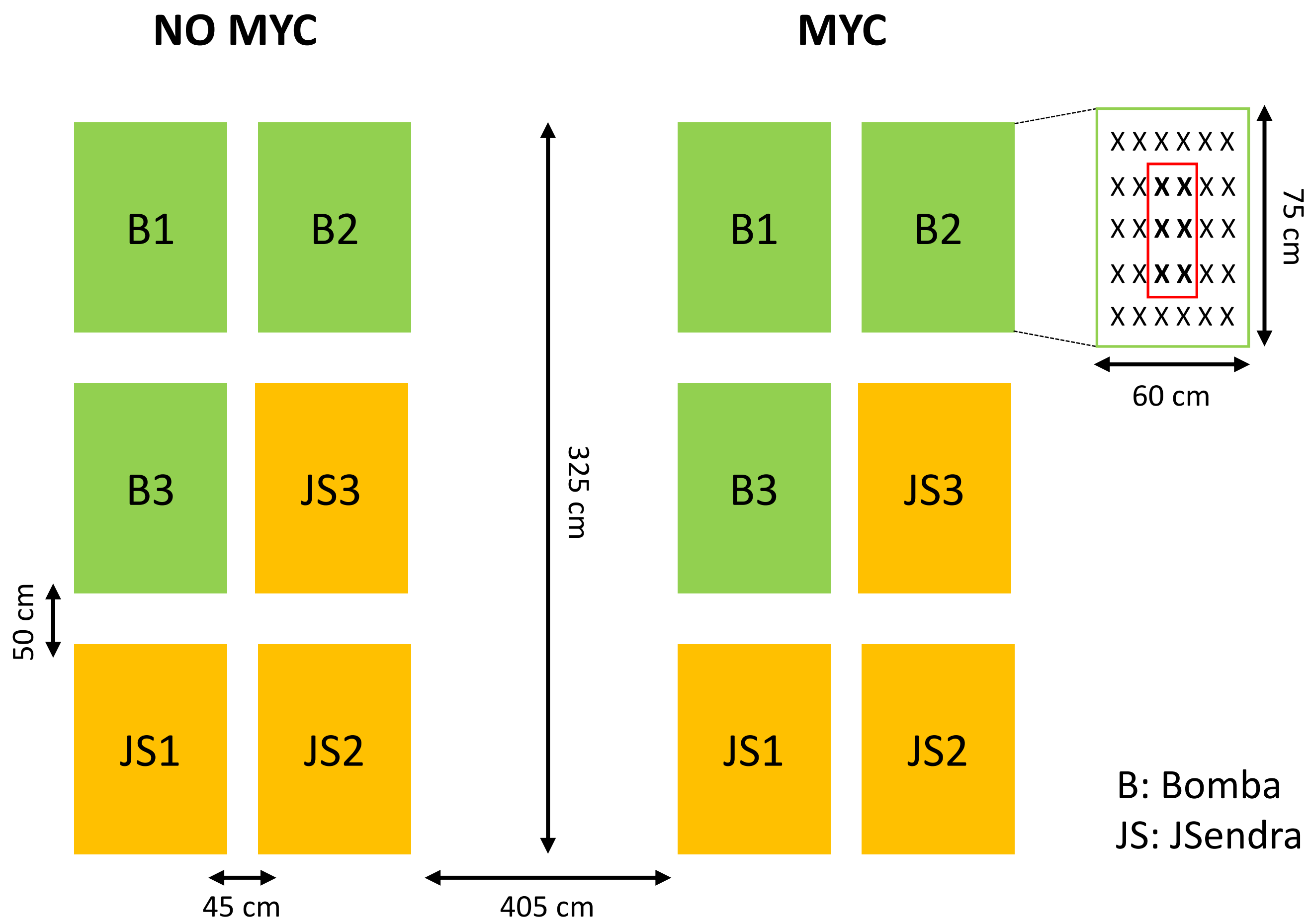


**Fig S1. Experimental design followed to grow rice plants in flooded rice fields**. Distribution of plots containing the rice varieties Bomba and JSendra, mock-inoculated and *R. irregularis* inoculated.

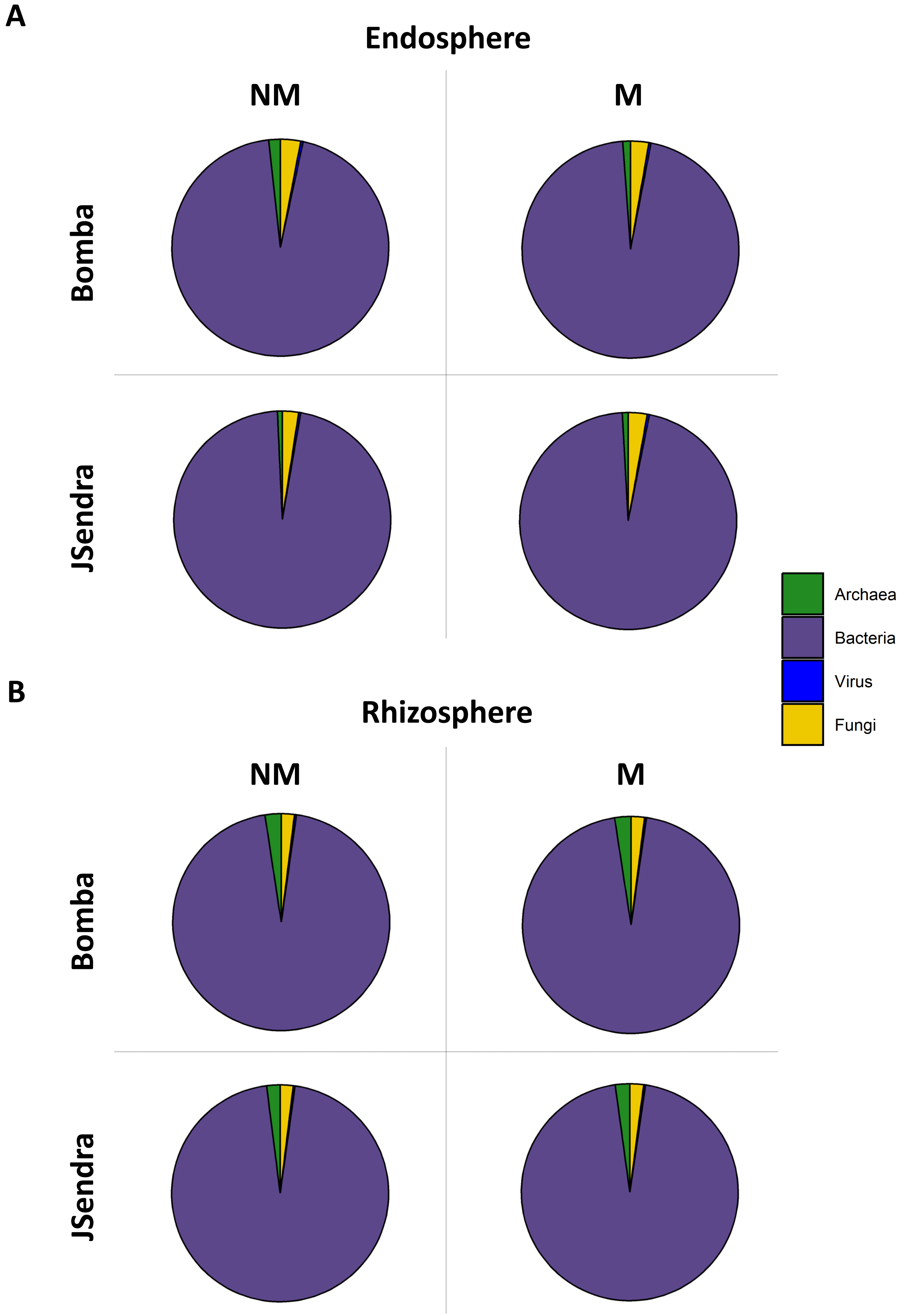


**Fig. S2. Proportion of reads assigned to Fungi, Bacteria, Archaea and Viruses in root endosphere and rhizosphere**. Bacteria, 94-96%, Fungi, 2-3%, Archaea, 1-2%, Viruses, 0.3%.

### Non – Mycorrhizal conditions

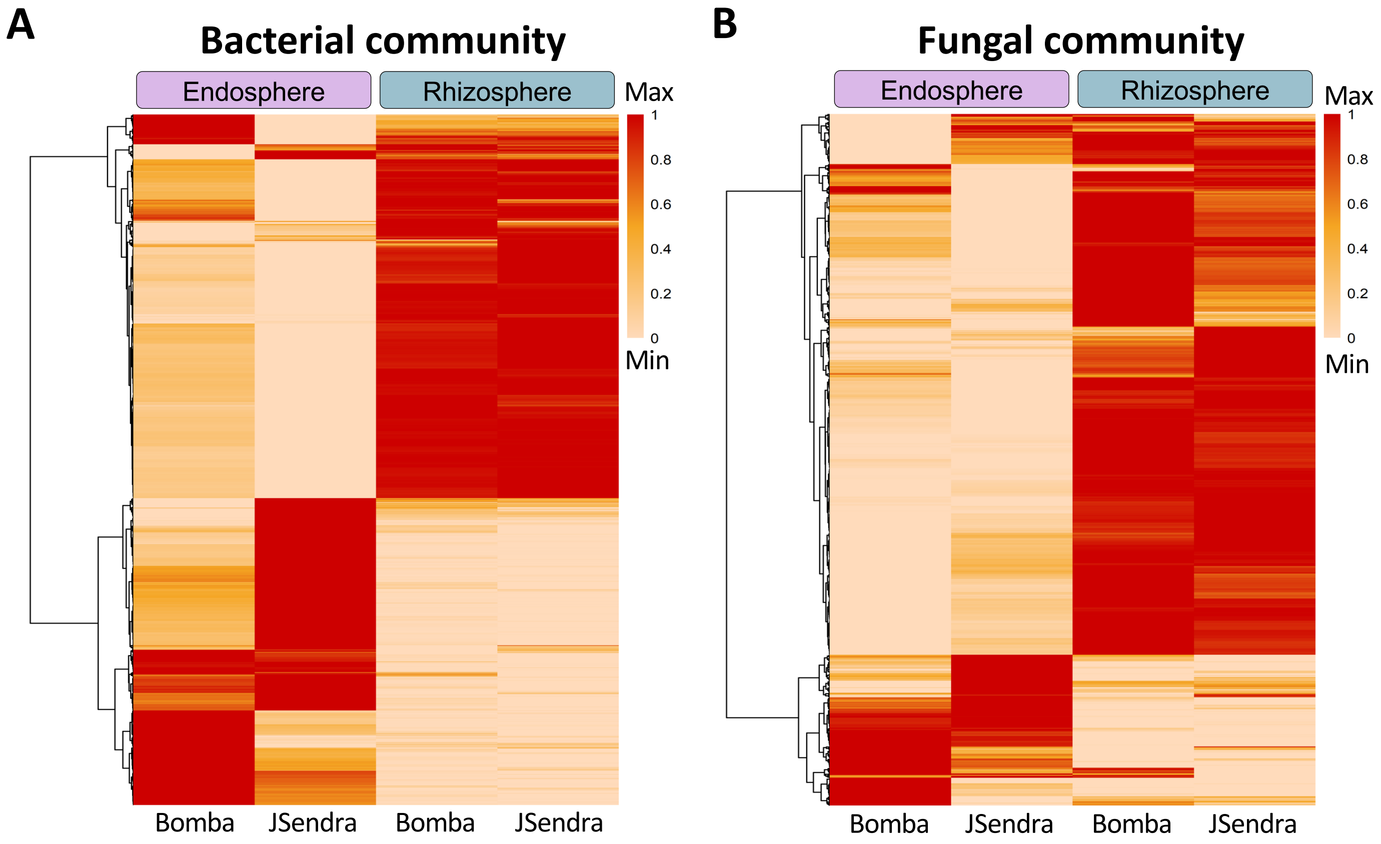


**Fig. S3. Differential abundance of microbial taxa in the root endosphere and rhizosphere of non-mycorrhizal Bomba and JSendra plants**. Data is presented at the species level. Heatmap represents bacterial and fungal species with a relative abundance greater than 0.01% in each sample. The Min-Max scaling normalization method was applied to the data after performing the $\log_2(x+1)$ transformation, rescaling it to the 0 to 1 range. The colour code indicates differences in the relative abundance ranging from light (less abundant) to dark (more abundant) red.

### Rhizosphere - Fungi

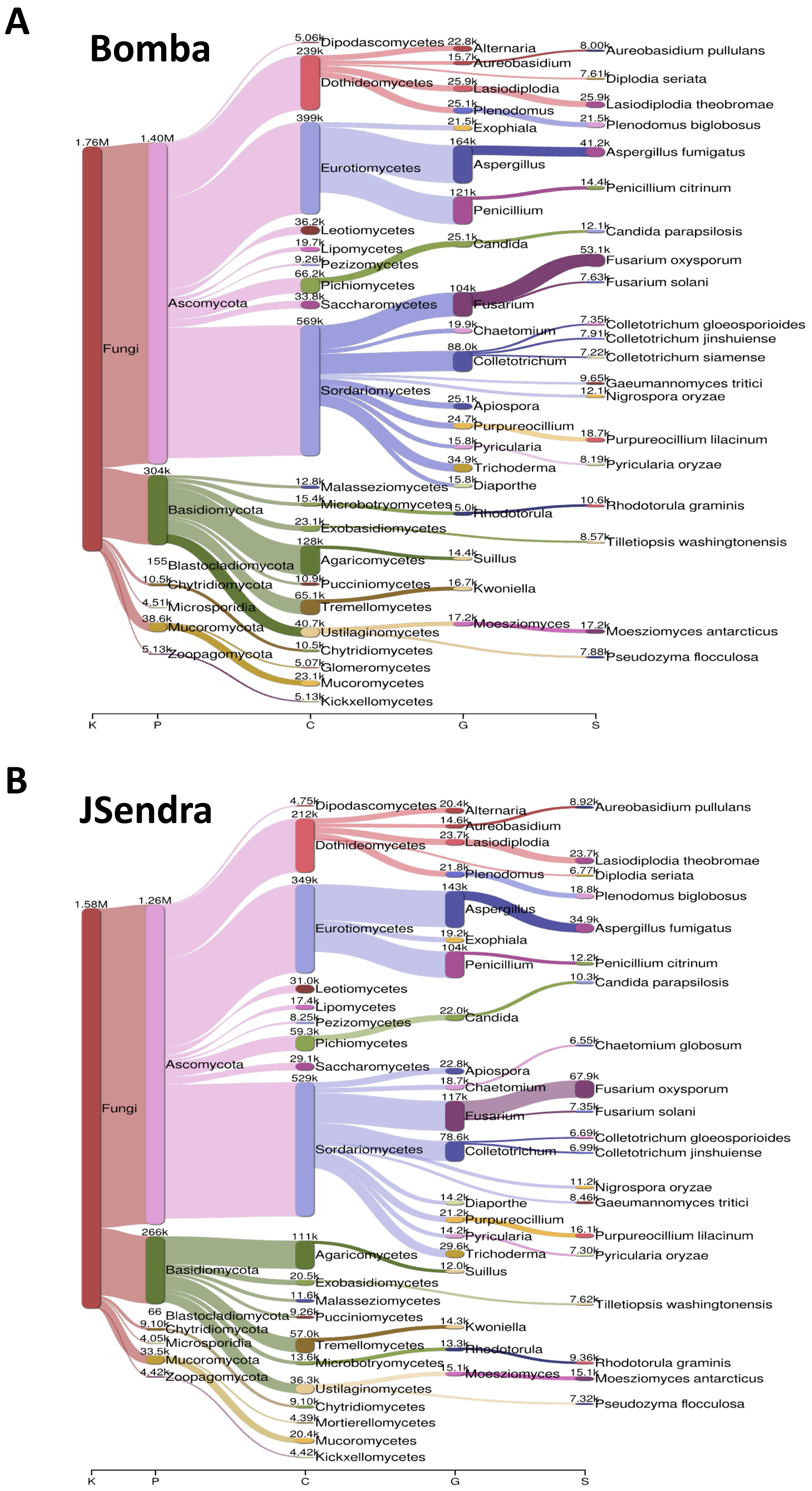


**Fig. S4. The most abundant fungal taxa identified in the rhizosphere of Bomba (A) and JSendra (B) plants**. Sankey diagrams showing the relative abundance of the top 20 taxa in each variety is presented, from the kingdom (on the left) to species (on the right). The width of the bar is proportional to the abundance, whereas nodes depict the hierarchy levels of microbial taxa.

### Mycorrhizal conditions

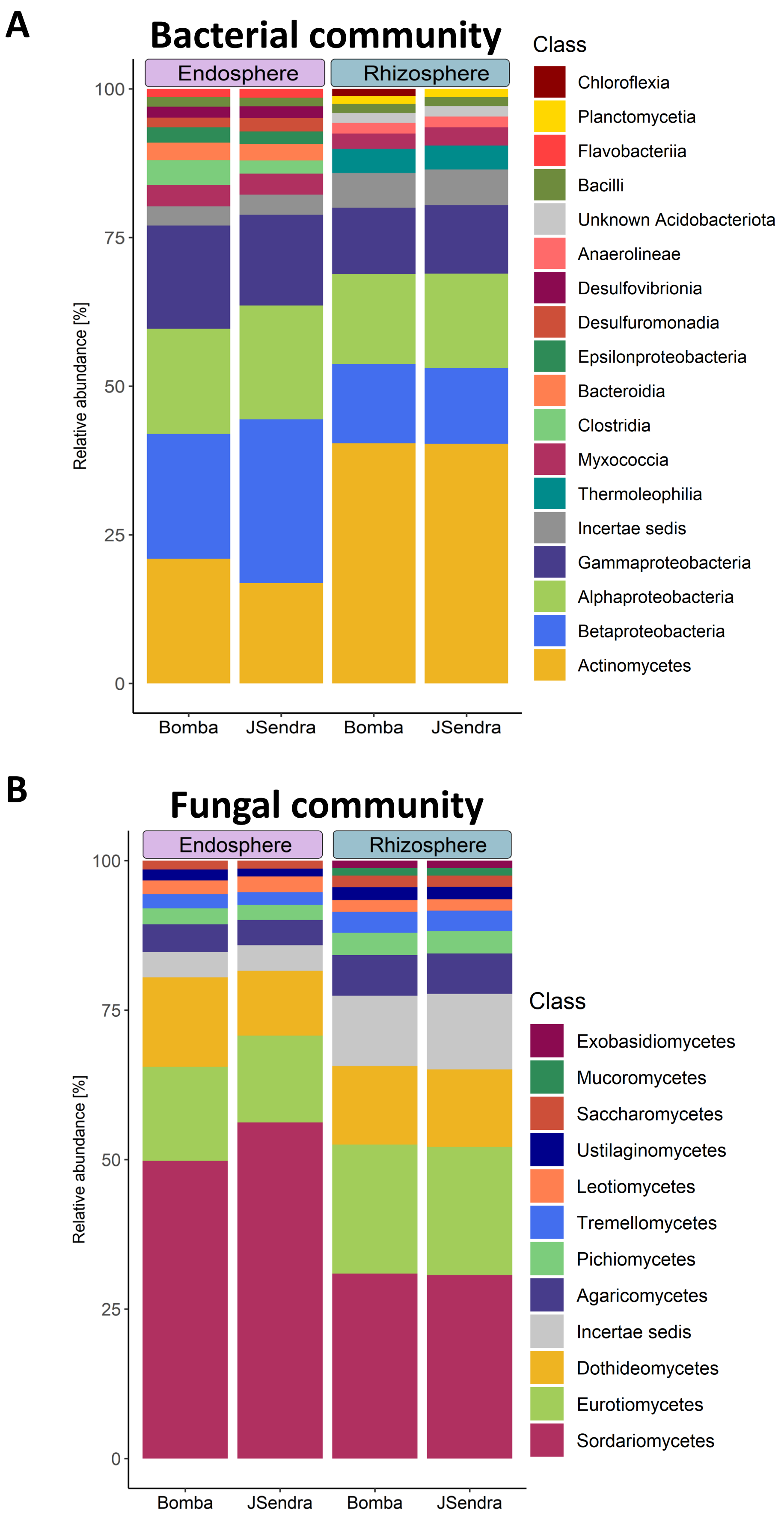


**Fig. S5. Taxonomic composition of the bacterial and fungal community in mycorrhizal roots of Bomba and JSendra plants**. Different colors indicate different taca at the Class level.

### Mycorrhizal conditions

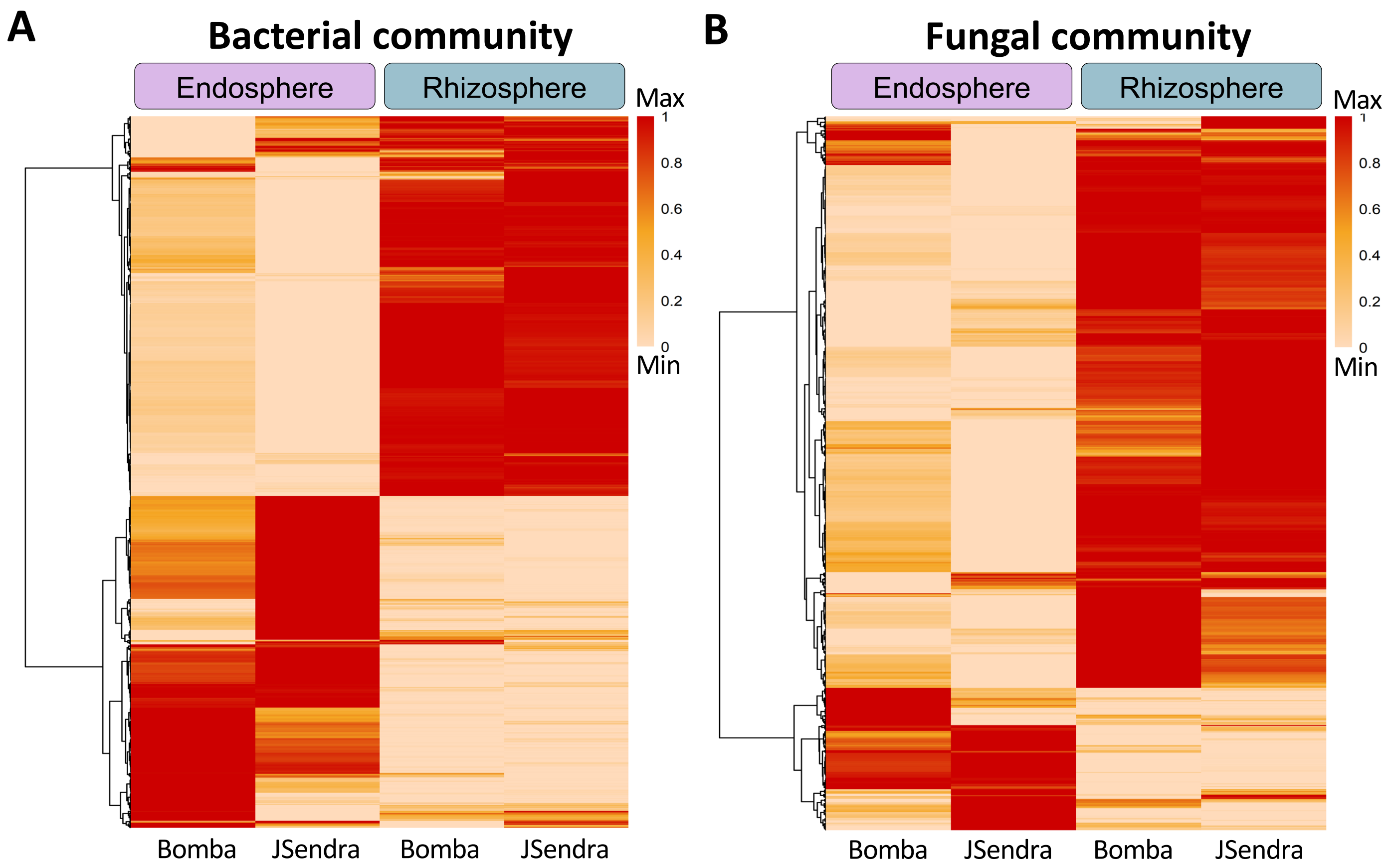


**Fig. S6. Differential abundance of bacterial and fungal taxa in the root endosphere and rhizosphere of mycorrhizal Bomba and JSendra plants**. Different colors indicate different taxa at the species level. Heatmap represents species with a relative abundance greater than 0.01% in each sample. The Min-Max scaling normalization method was applied to the data after performing the $\log_2(x + 1)$ transformation, rescaling it to the 0 to 1 range. The colour code indicates differences in the relative abundance ranging from light (less abundant) to dark (more abundant) red.

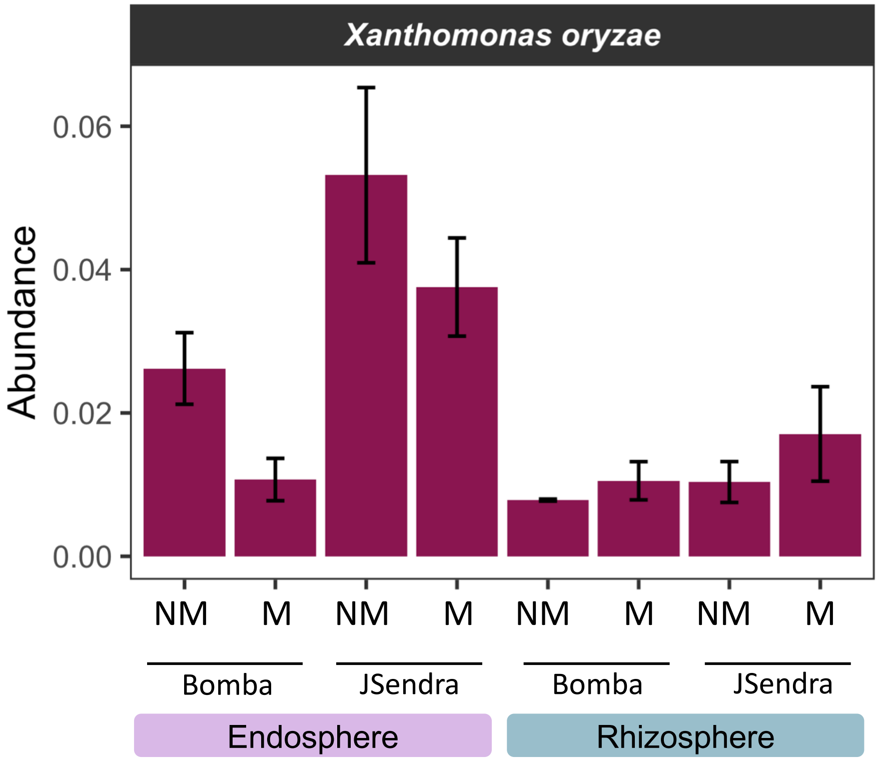


**Fig. S7. Relative abundance of *Xanthomonas oryzae*.** Data for non-mycorrhized and mycorrhized conditions from the endosphere and rhizosphere of Bomba and JSendra is shown.
